# Synergistic phenotypic shifts during domestication promote plankton-to-biofilm transition in purple sulfur bacterium *Chromatium okenii*

**DOI:** 10.1101/2023.10.20.563228

**Authors:** Francesco Di Nezio, Irvine Lian Hao Ong, René Riedel, Arkajyoti Goshal, Jayabrata Dhar, Samuele Roman, Nicola Storelli, Anupam Sengupta

## Abstract

The ability to isolate microorganisms from natural environments to pure cultures under optimized laboratory settings has markedly improved our understanding of microbial ecology. Laboratory-induced artificial growth conditions often diverge from those in natural ecosystems, forcing wild isolates into selective pressures which are distinct compared to those in nature. Consequently, fresh isolates undergo diverse eco-physiological adaptations mediated by modification of key phenotypic traits. For motile microorganisms, we still lack a biophysical understanding of the relevant traits which emerge during domestication, and possible mechanistic interrelations between them which could ultimately drive short-to-long term microbial adaptation under laboratory conditions. Here, using microfluidics, atomic force microscopy (AFM), quantitative imaging, and mathematical modelling, we study phenotypic adaptation of natural isolates of *Chromatium okenii*, a motile phototrophic purple sulfur bacterium (PSB) common to meromictic settings, grown under ecologically-relevant laboratory conditions over multiple generations. Our results indicate that the naturally planktonic *C. okenii* populations leverage synergistic shifts in cell-surface adhesive interactions, together with changes in their cell morphology, mass density, and distribution of intracellular sulfur globules, to supress their swimming traits, ultimately switching to a sessile lifeform under laboratory conditions. A computational model of cell mechanics confirms the role of the synergistic phenotypic shifts in suppressing the planktonic lifeform. Over longer domestication periods (∼10 generations), the switch from planktonic to sessile lifeform is driven by loss of flagella and enhanced adhesion. By investigating key phenotypic traits across different physiological stages of lab-grown *C. okenii*, we uncover a progressive loss of motility via synergistic phenotypic shifts during the early stages of domestication, which is followed by concomitant deflagellation and enhanced surface attachment that ultimately drive the transition of motile sulphur bacteria to a sessile biofilm state. Our results establish a mechanistic link between suppression of motility and surface attachment via synergistic phenotypic changes, underscoring the emergence of adaptive fitness under felicitous laboratory conditions that comes at a cost of lost ecophysiological traits tailored for natural environments.

**Graphical abstract:** 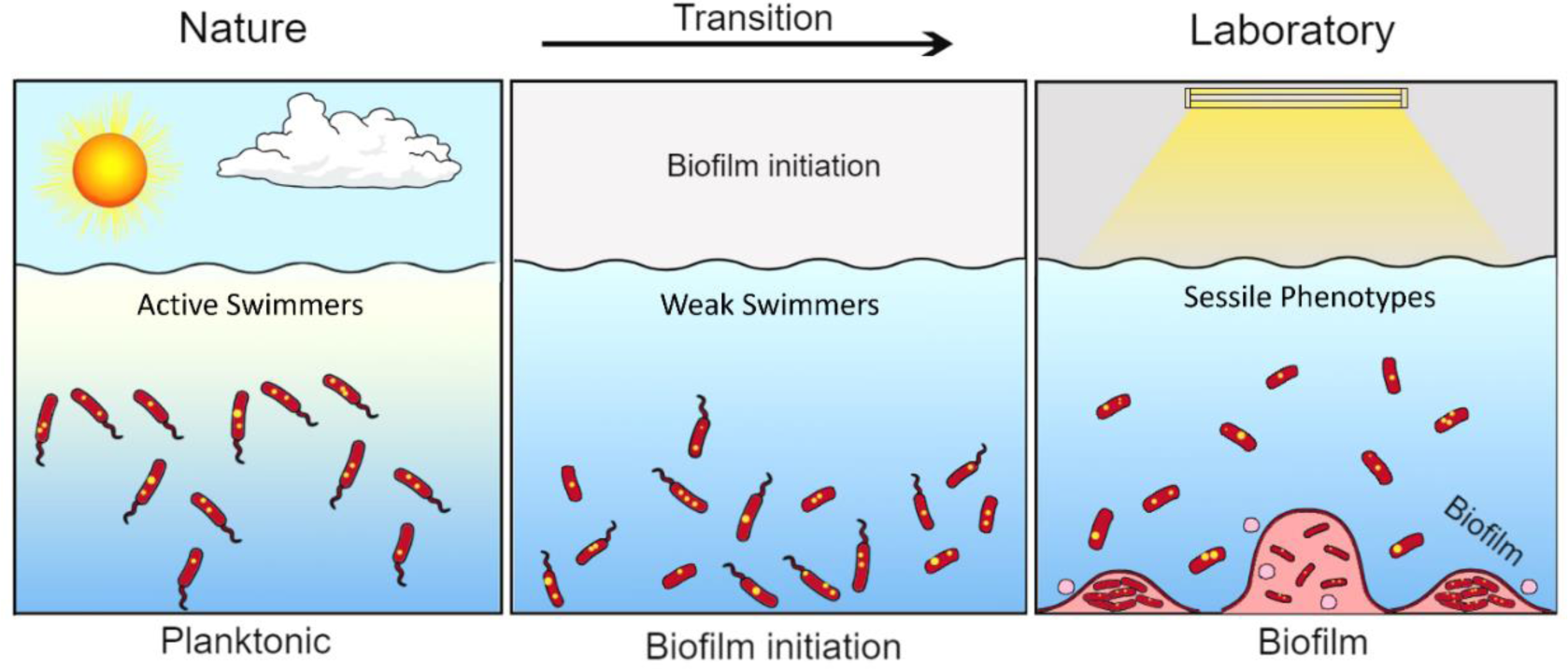

## Introduction

The ability to isolate microorganisms from natural settings and grow them under controlled laboratory conditions have driven our current understanding of the behaviour, physiology and fitness of microbes in a systematic manner. Natural microbial habitats offer conditions which are far from optimal, wherein diverse abiotic and biotic pressures shape microbial lifestyles and survival strategies (1, 2). A key distinction between the natural and lab-based conditions lies in the availability of nutrients, and other critical resources including secondary metabolites, which are central to optimal growth and fitness. Under the tailored growth conditions of laboratory settings, freshly isolated microorganisms experience a resource-replete setting, which often result in loss of key ecophysiological traits, following phenotypic adaptation or long-term mutations under favourable conditions (3–5), ultimately leading to selection of laboratory strains (1). Adaption to laboratory settings is characterized by changes in morphotype, physiology, and biological fitness, with the first signs of such diversification appearing within 2-3 days of domestication (3). The enhanced fitness under laboratory conditions comes at the cost of ecologically-relevant traits found otherwise in their naturally-occurring counterparts (6). The consequences of domestication vary widely across species, as well as within a domesticated population (3). While most studies to date have focused on sessile species, motile species growing in batch cultures may also show phenotypic alterations including loss of motility and associated rapid growth, possibly due to the higher costs of flagellar construction (7–9). Studies so far indicate that species may alter or lose multiple traits concomitantly (2, 7, 9), however little is known if the loss of a trait proceeds synergistically, or independent of other emerging traits. Despite the far-reaching implications of the nature-to-lab domestication among diverse microbial species, currently we lack a mechanistic understanding of the behavioural and physiological changes in relation to the emergent traits, and their co-evolution across different domestication timescales.

With an aim to bridge the current gaps in our understanding of microbial domestication and its implications, here we focus on the motile purple sulfur bacterium (PSB) *Chromatium okenii*, a member of the *Chromatiaceae* family which comprises physiologically similar species and genera of the γ-Proteobacteria, known to perform anoxygenic photosynthesis (10). PSB normally develop under anoxic conditions in the presence of light where sulfide (S^2-^) serves as an electron donor in the photosynthetic process and is oxidized to sulfate (SO_4_^2-^) through an intermediate accumulation of elemental sulfur (S^0^) within the cell (10, 11). The selective environmental factor that determines the development of PSB populations in aquatic ecosystems is the presence of a physical structure that prevents vertical mixing and allows the establishment of an anoxic compartment, such as meromictic lakes (12). The development of euxinic environments, with opposing gradients of oxygen and hydrogen sulfide (H_2_S), in the presence of light provides the ideal environment for the development of PSB, as well as the green sulfur bacteria (GSB) (13, 14). The euxinic conditions of Lake Cadagno, a meromictic lake located in the southern Swiss Alps, provide a conducive habitat for a thriving community of anoxygenic phototrophic sulfur bacteria (15). Within the distinctive bacterial layer of this lake, seven PSB species, including *C. okenii*, and two GSB species are discernible, and they play pivotal roles in the lake’s major biogeochemical processes (16–18). Over the years, it has been possible to isolate and cultivate in the laboratory all nine species present in the bacterial layer of Lake Cadagno (19–21). However, cultivation of microorganisms in laboratory settings involves the use of highly nutrient-rich growth media that deviate from the natural environment from which they are isolated (1).

PSB *C. okenii* is a positively phototactic and negatively aerotactic species (22), and its flagellar motility provides it with a distinct advantage, through the process of bioconvection (23), allowing it to reach the most favorable environmental niches and compete effectively with other microbes (Di Nezio *et al*., under review). However, maintaining a flagellar motility system is energetically costly and, given the necessity of this complicated system for bacterial survival, its regulatory efficiency is under considerable selective pressure in the environment (24, 25). One of the main factors determining motility is cell morphology (26–28), which strictly depends on the environmental conditions bacterial cells face. When facing environmental challenges like antibiotics and predation, bacteria can gain advantages over their free-floating counterparts by forming biofilms, thereby increasing both cohesion among cells and adhesion to solid surfaces and transitioning to a non-motile state (29).

Natural habitats are characterized by variability, which can lead to persistent stress situations (e.g., nutrient depletion, chemical inhibition, temperature shifts). If organisms can reduce their exposure to such stressors, or are phenotypically prepared for anticipated changes before they occur, they may perform better and be more likely to persist (30). For instance, many microorganisms coping with fluctuating environments have developed the ability to store surplus of various substances during unstable growth. PSB are known for their ability to store reserve substances within their cells, such as glycogen, polyhydroxybutyrate (PHB), and sulfur globules (SGBs) (31). Sulfur globules (SGBs), stored intracellularly by PSB, can be further oxidized into sulfate when the availability of reduced sulfur compounds necessary for anoxygenic photosynthesis is scarce (11, 32).

In general, little is known about the mechanisms, extent and the rate at which physiological and behavioural traits alter, during domesticating wild microorganisms. More specifically, there is a significant knowledge gap concerning the domestication processes that microorganisms dwelling in extreme habitats, such as *C. okenii*, undergo. Here, we use a combination of microfluidics, atomic force microscopy (AFM), quantitative imaging, and mathematical modelling to study biophysical changes in natural isolates of *C. okenii* as they grow over multiple generations under suitable laboratory conditions. We report concomitant changes in multiple phenotypic traits which, acting in a synergistic manner, supress the motility during the nature-to-lab domestication phase. Within timescales of 8-10 generations, the populations switched from a planktonic to sessile biofilm lifeform, mediated by high cell-surface adhesion and loss of flagella. We present below the observed changes and develop a cell-based mechanistic model to delineate the impacts of domestication, and discuss their physiological implications on the fitness of lab-grown *C. okenii*

## Results

### Domestication modifies cell morphology and intracellular SGBs attributes

To uncover the reason behind the differences observed in *C. okenii* motility between wild and domesticated populations, we looked for potential alterations of the morphological features of the cells. Cell phenotype and SGBs characteristics were monitored in the two different artificial growing conditions of the window-sill (WND) and the incubator (INC), and compared with cells freshly isolated from Lake Cadagno (Lake), using cell-level quantitative imaging (Table 1). In laboratory-grown cultures, INC cells displayed a higher growth rate, 0.0081 ± 0.0031 h^-1^ (doubling time ∼86 ± 27.1 h), compared to the WND population with a growth rate of 0.0045 ± 0.0022 h^-1^(doubling time ∼156.1 ± 52.9 h, Figure 1a). During the course of population growth, the aspect ratio of cells (length / width) increased, reaching its maximum in the exponential phase, under both laboratory growing conditions compared to natural samples (Figure 1b), in which cells retained a more rounded shape (lower aspect ratio).

**Figure 1.**
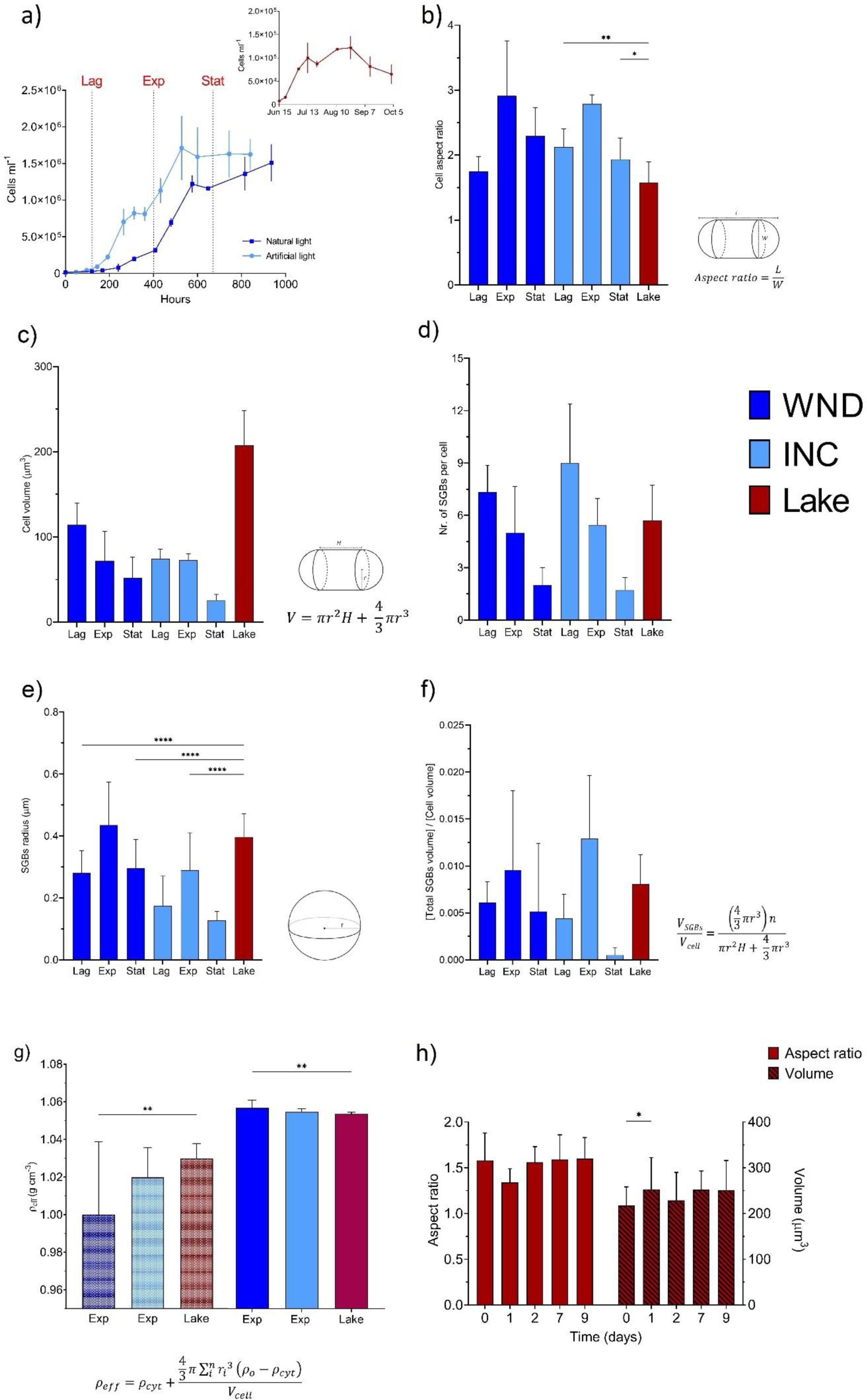
Domestication alters cell morphology and SGB characteristics. **a)** Growth curves of domesticated cells under the two laboratory conditions tested. Inlet shows number *C. okenii* cells in the lake across the whole 2022 sampling season. **b)** Domesticated cells undergo a similar modification of the aspect ratio under artificial growing conditions. **c)** Reduction in the volume of laboratory-grown cells over different growth stages compared to lake-sampled cells volume. **d)** Decrease in the number of intracellular SGBs in different stages of growth. **e)** Variation of SGBs size (length of radius) and **f)** total SGBs / cell volume ratio occurring over cell growth. **g)** Differences in densities of *C. okenii* cells in the absence (speckled colored bars) and presence (full colored bars). One-way ANOVA, *P* < 0.01; post hoc Dunnet test; asterisks indicate statistically significant difference. Error bars represent standard deviation (N=20). **h)** Wild *C. okenii* cells exhibit consistency in the main morphological traits across temperature variations (4°C to 20°C) from natural to laboratory environments, as evidenced by aspect ratio and volume measurements. 2-way ANOVA, *P* < 0.01; post hoc Tukey’s test. Error bars represent standard deviation (N=20).

**Table 1.** Mean values ± SD (in μm) of cell shape and volume, SGBs number, size and total volume per cell for *C.okenii*.

| Growth stage |  | Cell morphology |  |  | SGBs |  |  |
| --- | --- | --- | --- | --- | --- | --- | --- |
| | | Length | Width | Volume | Number | Radius | $V_{\text{SGBs}}/V_{\text{cell}}$ |
| WND | Lag | $8.95 \pm 0.93$ | $4.58 \pm 0.27$ | $114.35 \pm 25.32$ | $7 \pm 1$ | $0.28 \pm 0.07$ | $0.006 \pm 0.002$ |
| | Exponential | $8.36 \pm 0.77$ | $4.16 \pm 0.85$ | $71.75 \pm 34.63$ | $5 \pm 3$ | $0.43 \pm 0.13$ | $0.009 \pm 0.008$ |
| | Stationary | $7.91 \pm 1.17$ | $3.75 \pm 0.63$ | $52.54 \pm 24.16$ | $2 \pm 1$ | $0.29 \pm 0.09$ | $0.005 \pm 0.007$ |
| INC | Lag | $8.34 \pm 1.08$ | $3.50 \pm 0.21$ | $72.94 \pm 7.41$ | $9 \pm 3$ | $0.16 \pm 0.07$ | $0.004 \pm 0.002$ |
| | Exponential | $9.36 \pm 0.51$ | $3.35 \pm 0.10$ | $69.90 \pm 12.47$ | $5 \pm 1$ | $0.29 \pm 0.12$ | $0.013 \pm 0.006$ |
| | Stationary | $5.25 \pm 0.90$ | $2.72 \pm 0.23$ | $25.51 \pm 7.20$ | $2 \pm 0$ | $0.13 \pm 0.02$ | $0.001 \pm 0.0001$ |
| Lake | Exponential | $9.91 \pm 1.73$ | $6.33 \pm 0.38$ | $207.66 \pm 40.70$ | $6 \pm 2$ | $0.40 \pm 0.08$ | $0.008 \pm 0.003$ |

Cell volume also increased, reaching a maximum at the lag growth stage in both laboratory settings (114 ± 25 and 72 ± 7 μm^3^ at the stationary phase, WND and INC, respectively), while lake-sampled cells showed to be nearly 2-fold larger (207 ± 40 μm^3^, Figure 1c). Thereafter the cell size reduced as the population entered the late stationary stage (t > 700 h). Correspondingly, the number of the SGBs decreased over growth while their size (radius length) increased from lag to exponential phase, with INC cells showing a higher number of sulfur globules, although smaller in size (Figure 1d, e). The peak in SGBs size observed in the exponential phase of both domesticated populations is corroborated by the globules volume relative to the cell size (total globules volume/cell volume) shown in Figure 1f, which significantly enhanced when the populations entered the exponential growth stage (ANOVA: *P* < 0.01). During the transition to the stationary growth stage (600 h < t < 800 h), SGBs decreased by nearly 50% and 83%, in the WND and INC cells, respectively (Figure 1f).

### Effective cellular mass density

The results in Figure 1g show how variations in the specific content of SGBs affects the cell density. Particularly, values of cell effective density were significantly higher in the presence of intracellular SGBs than when the density of the SGBs was subtracted from the overall cell density, the difference being the density of the structural cell material. Such difference was more pronounced in laboratory cells at their exponential phase under both artificial growth conditions, when SGBs size and the ratio of globules volume over cell volume reached their max (Figure 1e, f). The presence of SGBs resulted an increase in cell density from 1.000 to 1.055 for WND, and from 1.019 to 1.053 g cm^-3^ for the INC populations (Figure 1g). Overall, the effective cellular mass density for the lab-grown cells were 0.3 % higher, in comparison to that of the cells from the lake (1.052 g cm^-3^).

### Loss of flagella

Alongside variations of the cellular morphology, mass density and SGBs attributes, we recorded a gradual loss of flagella in the INC lab-grown *C. okenii* population. As shown in Figure 2a, SEM and phase contrast micrographs track the presence of a polar flagella bundle on the cells freshly sampled from the lake chemocline. In contrast, cells lacking flagella were detected in the domesticated samples (Figure 2b, right), even though the imaging conditions were the same for both samples. Overall, close to 75% of cells grown under laboratory conditions showed loss of flagella by the time the population reached exponential phase, which further increased to ∼100% by the time the population reached stationary phase.

**Figure 2.**
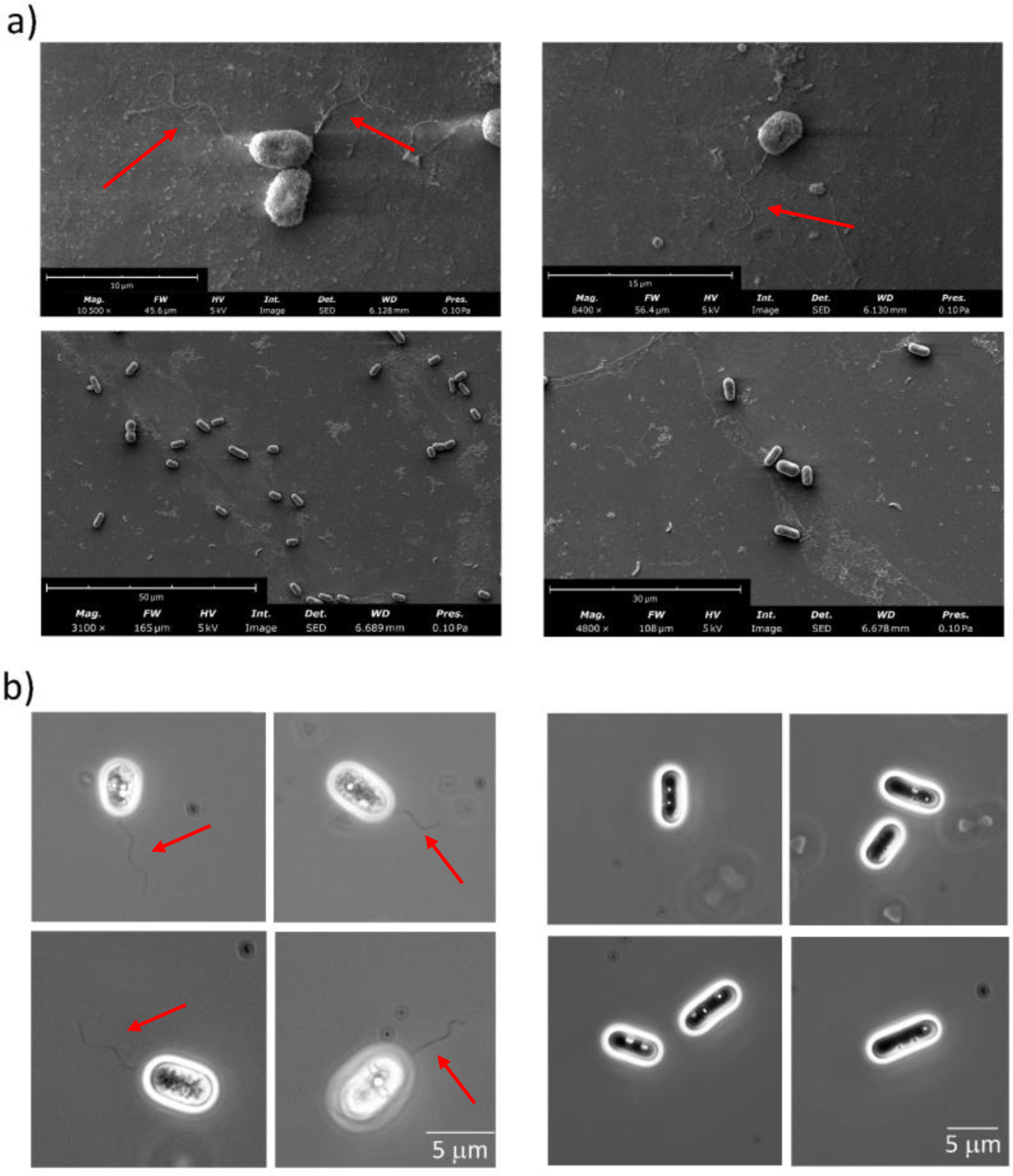
*Chromatium okenii* lose flagella during domestication. **a)** Scanning electron microscope images of *C. okenii* cells freshly sampled from the lake (upper panel) and laboratory-grown cells (lower panel). Red arrows indicate the polar flagellar tuft. **b)** Microscopy images (100X, phase contrast) of *C. okenii* cells from a fresh lake water sample (left panel) and INC laboratory-grown cells right panel). No flagella are visible in domesticated cells. Intracellular sulfur globules are visible as highly refractive spheres. Red arrows indicate the polar flagellar tuft.

### Emergence of planktonic to surface-associated lifestyle under laboratory growth conditions

PSB *C. okenii* cells coming from vastly different conditions, wild (freshly sampled from the lake) and domesticated (laboratory-grown) (Table 1), were compared to investigate how adaptation to artificial settings shapes cell phenotypic traits, amenable to their physiological growth stage. At first, we tracked the motility of both lake-sampled and domesticated *C. okenii*. We observed a change in the swimming speed between lake and laboratory populations, and between WND and INC cells as well (Figure 4b and c). For the WND population, *C. okenii* swims at 11.84 ± 2.73 μm s^-1^ during the stationary stage, in contrast to the swimming speed during the exponential stage ranging around 12.63 ± 1.96 μm s^-1^, as shown in Figure S1. In INC cells, the swimming speed during the early stationary phase was 6.59 ± 4.27 μm s^-1^, while for cells in exponential phase, it was 3.76 ± 0.63 μm s^-1^. Swimming speed for lake-sampled cells in exponential stage was 19.25 ± 1.86 μm s^-1^.

**Figure 3.**
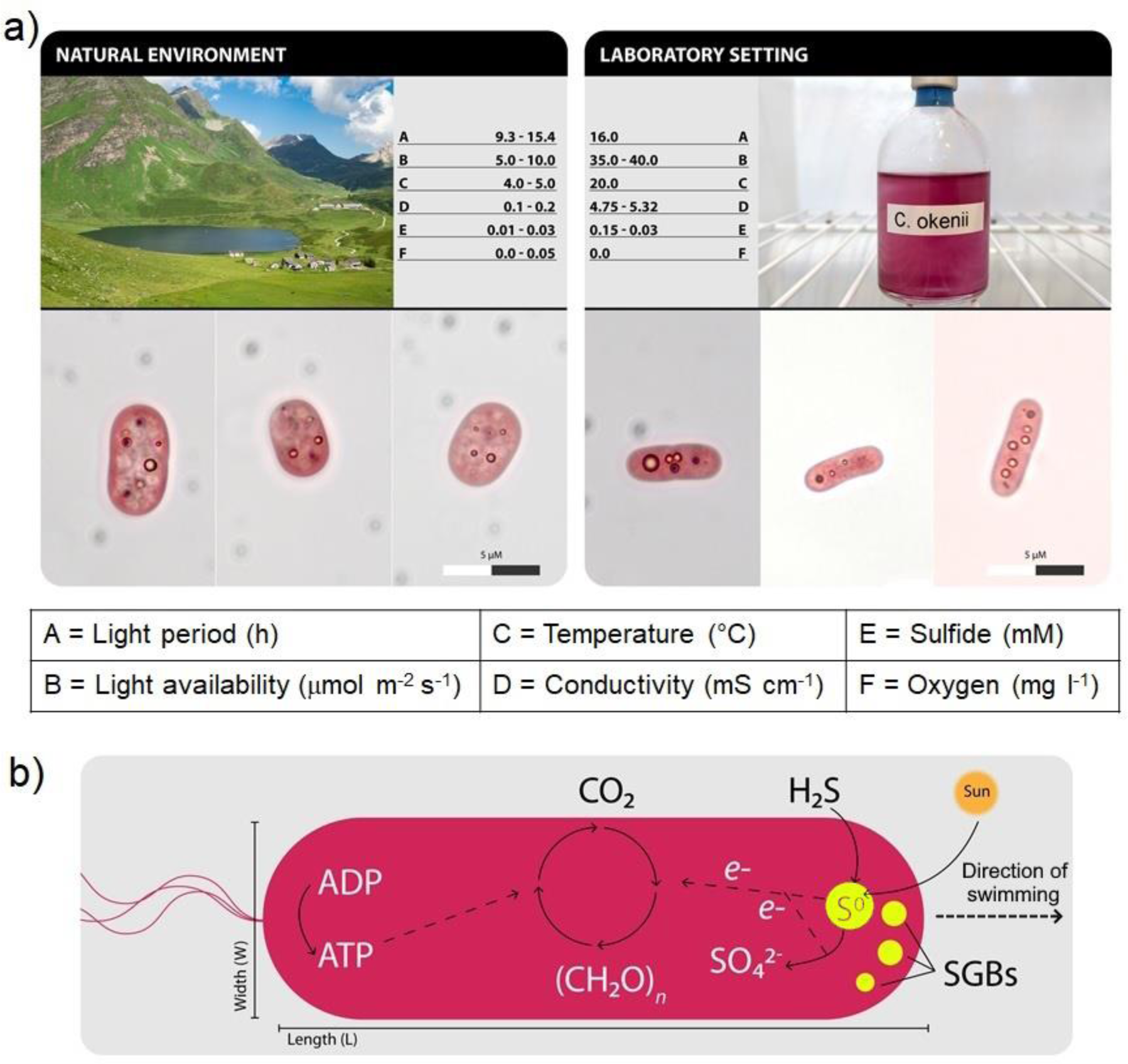
Difference in the physicochemical parameters measured between natural and artificial environments. **a)** *Upper half* – Different values of the main abiotic factors (A-F) influencing the growth of *C. okenii* in the natural and laboratory environment; numbers represent seasonal ranges. *Lower half –* light photomicrographs showing morphological differences between wild and domesticated *C. okenii* cells. **b)** Schematics of energy and reducing power synthesis in anoxygenic phototrophs. Yellow circles represent the sulfur globules inside the cells produced from the oxidation of H_2_S. Length, width and SGBs number are the main features used to characterize cell morphology.

**Figure 4.**
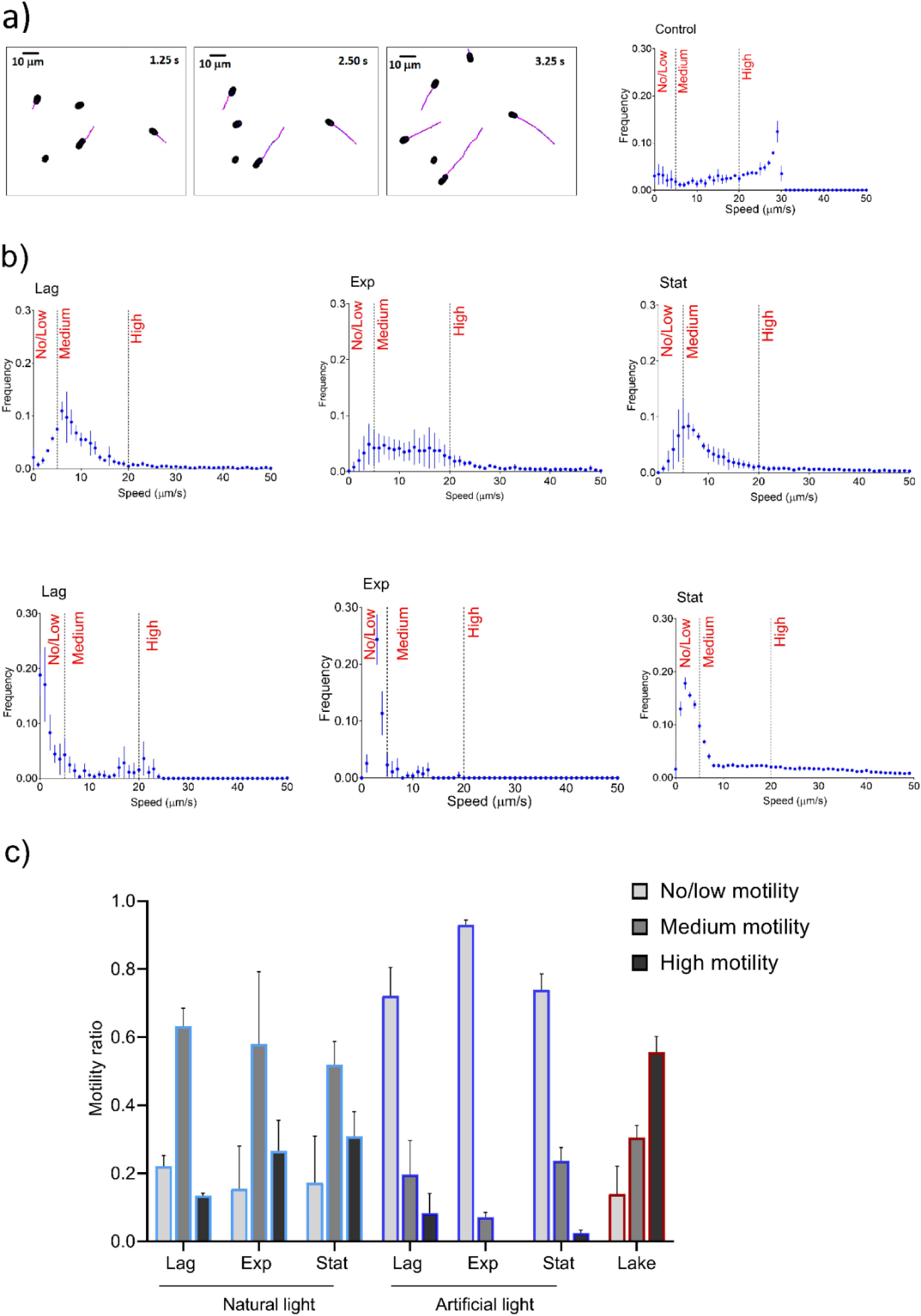
Lake-sampled cells display a higher motility than domesticated cells. **a)** Segmented images of lake-sampled cell trajectories over a 3.75 s acquisition and their relative speed distribution (control). **b)** Histograms of the distribution of speeds of laboratory-grown cells in lag, exponential and stationary phase under two artificial growing conditions (WND top histograms and INC bottom histograms). **c)** Bar plot of the different ratios of domesticated and natural cells within the three motility regimes. Error bars represent standard deviation (N=3).

We divided the observed motility into three regimes, no/low motility (< 5 μm s^-1^), medium motility (5 - 20 μm s^-1^), and high motility (> 20 μm s^-1^). Bar plot in Figure 4c shows cells distribution among the three motility regimes. The wild population showed higher motility compared to both domesticated cultures, with 55% of the lake-sampled cells falling into the category of high motility (swimming speed > 20 μm s^-1^) compared to a value of 27% and 0% for the WND and INC populations in exponential phase, respectively. It is worth noting that, over longer time periods, emergence of a medium-motility subpopulation in cells grown under natural light on the window-sill, and a low-motility subpopulation in cultures grown under artificial light in the incubator (Figure 4c) was recorded, suggesting a gradual suppression of the planktonic behavior in favour of a surface-associated lifestyle with an increasing degree of domestication. Furthermore, the natural light conditions, as compared to the artificial light conditions in incubator promote higher motility. Evidence of the key role played by light, particularly in terms of nature and duration of the light period for the ecophysiology of *C. okenii* comes from different growth rates observed in WND and INC cells, with light/dark photoperiods of 12/12 h and 16/8 h, respectively (Figure 1a). The numerical ratios quantifying the mobility of highly motile cells in contrast to cells displaying moderate and low motility indicate the variability in motility exhibited by domesticated populations throughout their growth under distinct laboratory settings (Figure 4). The motility of WND cells consistently surpasses that of INC cells, with these observed ratios significantly declining in comparison to those observed within the lake population (Figure 4a, b). Furthermore, the assessment of mean velocity across the three experimental conditions substantiates these findings (Figure 4c).

### Light availability impacts motility of *C. okenii*

To further investigate the role of domestication in determining alterations of phenotypic traits, cells were tested for their phototactic response, a key trait of *C. okenii* wild population (Figure 5). In general, we observed that light triggered a phototactic response in *C. okenii* wild cells after a period of incubation in the dark (1 h), with a swimming speed that increased after exposure to the light source. On the contrary, laboratory-grown cells showed almost no reaction to light exposure (Figure 6). Lake-sampled cells kept in the dark after 30 and 90 minutes were used as a control.

**Figure 5.**
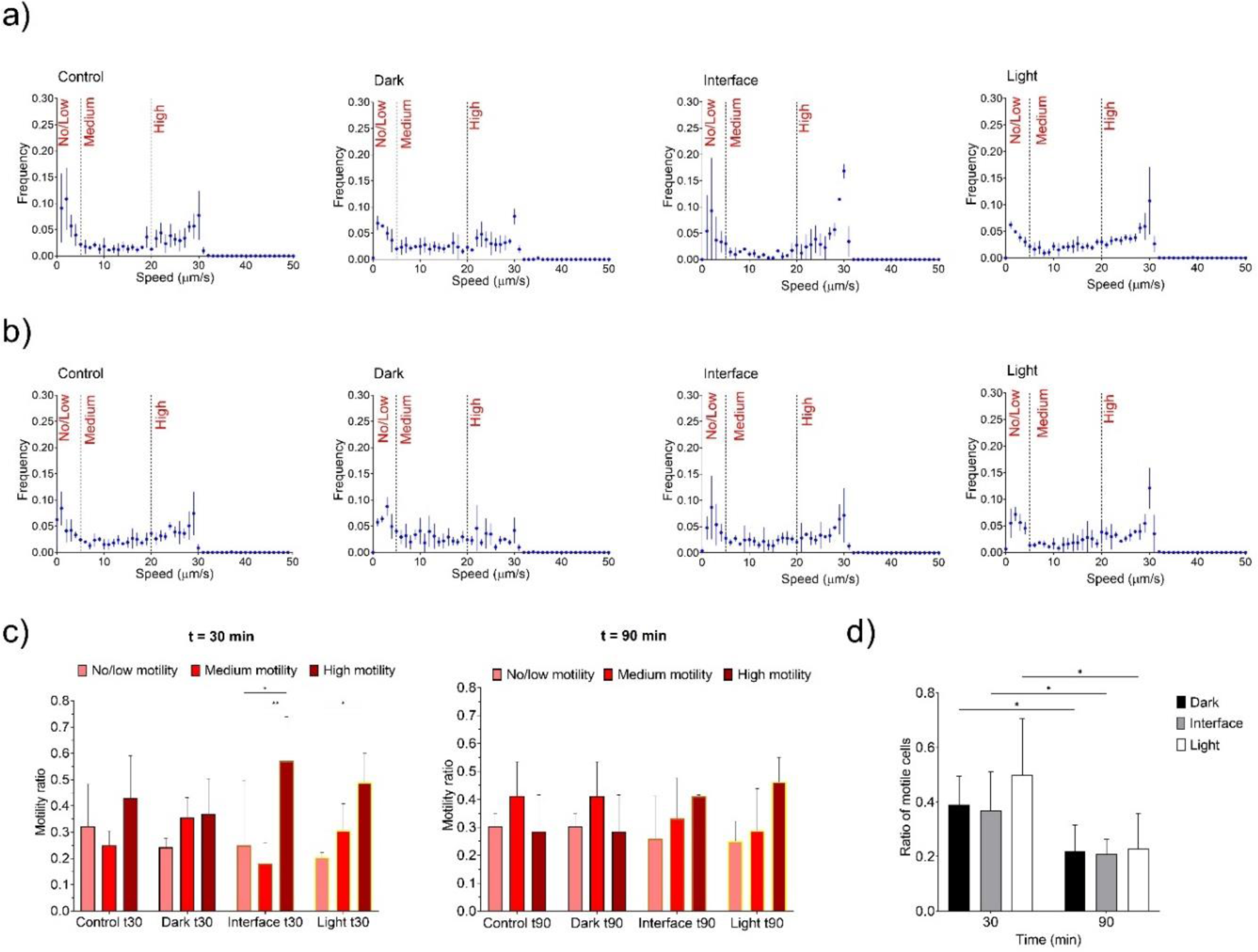
Lake-sampled cells display phototactic behavior and higher motility than domesticated cells. Different swimming speed distribution of lake-sampled cells after 30 (**a**) and 90 (**b**) min of light exposure in a half-shaded, half-illuminated microfluidic chip (see Figure S3). Black, dashed lines indicate the three different motility regimes chosen (see Materials and Methods). **c)** Bar plots show distribution of cells within the three motility ranges. **d)** Ratio of motile *vs* non-motile cells for both phototaxis experiment. Two-way ANOVA, *P* < 0.01; post hoc Tukey test; asterisks indicate statistically significant differences. Error bars represent standard deviation (N=3).

**Figure 6.**
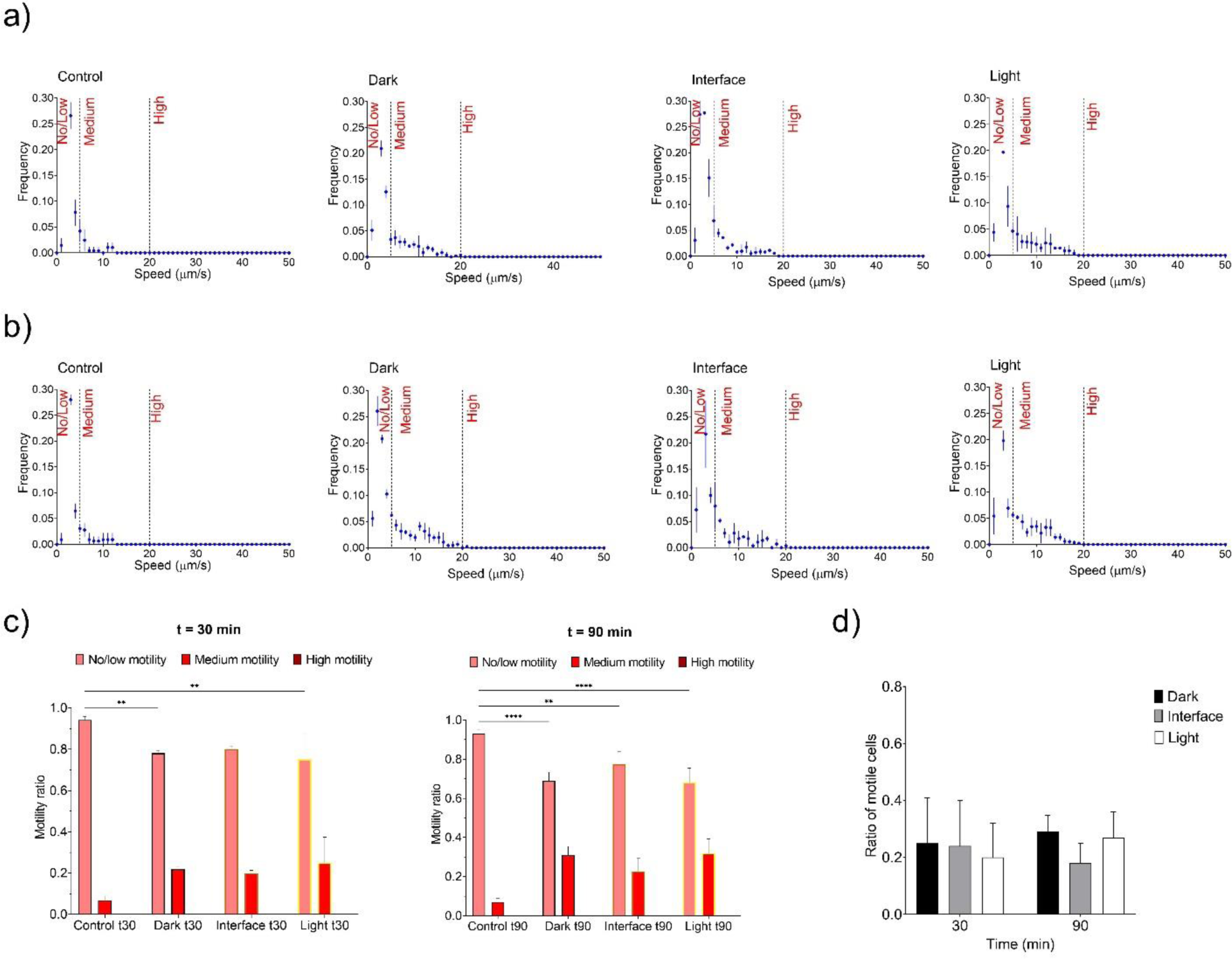
Laboratory-grown cells display no response to light and little motility. Swimming speed distribution of laboratory-grown cells after 30 (**a**) and 90 (**b**) min of light exposure in a half-shaded, half-illuminated microfluidic chip. Black, dashed lines indicate the three different motility regimes chosen (see Materials and Methods). **c)** Bar plots showing distribution of cells within the three motility ranges. **d)** Ratio of motile *vs* non-motile cells. Two-way ANOVA, *P* < 0.01; post hoc Tukey test; asterisks indicate statistically significant differences. Error bars represent standard deviation (N=3).

Wild cells exposed to light within a millifluidic confinement, half of which was covered to block incoming light (Figure S3), displayed a higher motility than cells located in the dark half and at the light-dark interface. After 30 min, we observed a higher frequency of speeds above 20 μm s^-1^ in cells exposed to light, compared to the control, a range of values that falls under the category we defined as ‘high motility’ (Figure 5a). On the contrary, the control and cells kept in the dark were characterised by almost uniform values for each speed regime, indicating that the motility did not change with light (Figure 5a, c left bar plot). After 90 min of light exposure, cells appeared to be less photo-responsive, as there was no significative difference between the three conditions, with similar ratios for each motility regime (Figure 5b, c right bar plot). Interestingly, we also observed notable differences in the way cells distributed throughout the millifluidic chamber (Figure S4). At *t_30_*, *C. okenii* cells were significantly more abundant in the illuminated half of the chamber, their number progressively decreasing towards the shaded half. According to the uniformity of the speed distribution observed, no significant differences were found in the cell distribution at *t_90_* (Figure S4a, b). These observations are also supported by the different ratios of motile *vs* non motile cells, revealing an overall larger fraction of motile cells at *t_30_* than at *t_90_* (Figure 5d). Similar phototactic behaviour was observed in previously dark incubated cells after 30 min of localized LED illumination at two different light intensities (Figure S5 and Supplementary Text 1).

Instead, domesticated cells displayed almost no response to light when loaded in the same millifluidic chip. Histograms of speed distribution at t_30_ and t_90_ of laboratory-grown *C. okenii* (Figure 6a, b) show how cell swimming activity remained unaffected between the illuminated and dark region of the millifluidic device. In fact, most of the domesticated cells fell within the ‘no/low motility’ regime, and none reached ‘high motility’ at both time points (Figure 6c). The absence of any significative difference in the motile *vs* non motile cell ratio, as well as in their distribution at both time points, across the three sections of the millifluidic device further confirms these observations (Figure 6d, S4b).

### Cell adhesion

We employed atomic force microscopy (AFM, see Materials and Methods) to measure the change in adhesive interactions of the lake and lab-grown *C. okenii* cells. As shown in Figure 7d, freshly isolated *C. okenii* cells had a cell-surface adhesion of 0.211 ± 0.091 nN (maroon boxplot), whereas after ∼8 generations of domestication, the cell-surface adhesion enhanced significantly, by ∼ 4-fold to 0.836 ± 0.584 nN (light blue boxplot).

**Figure 7.**
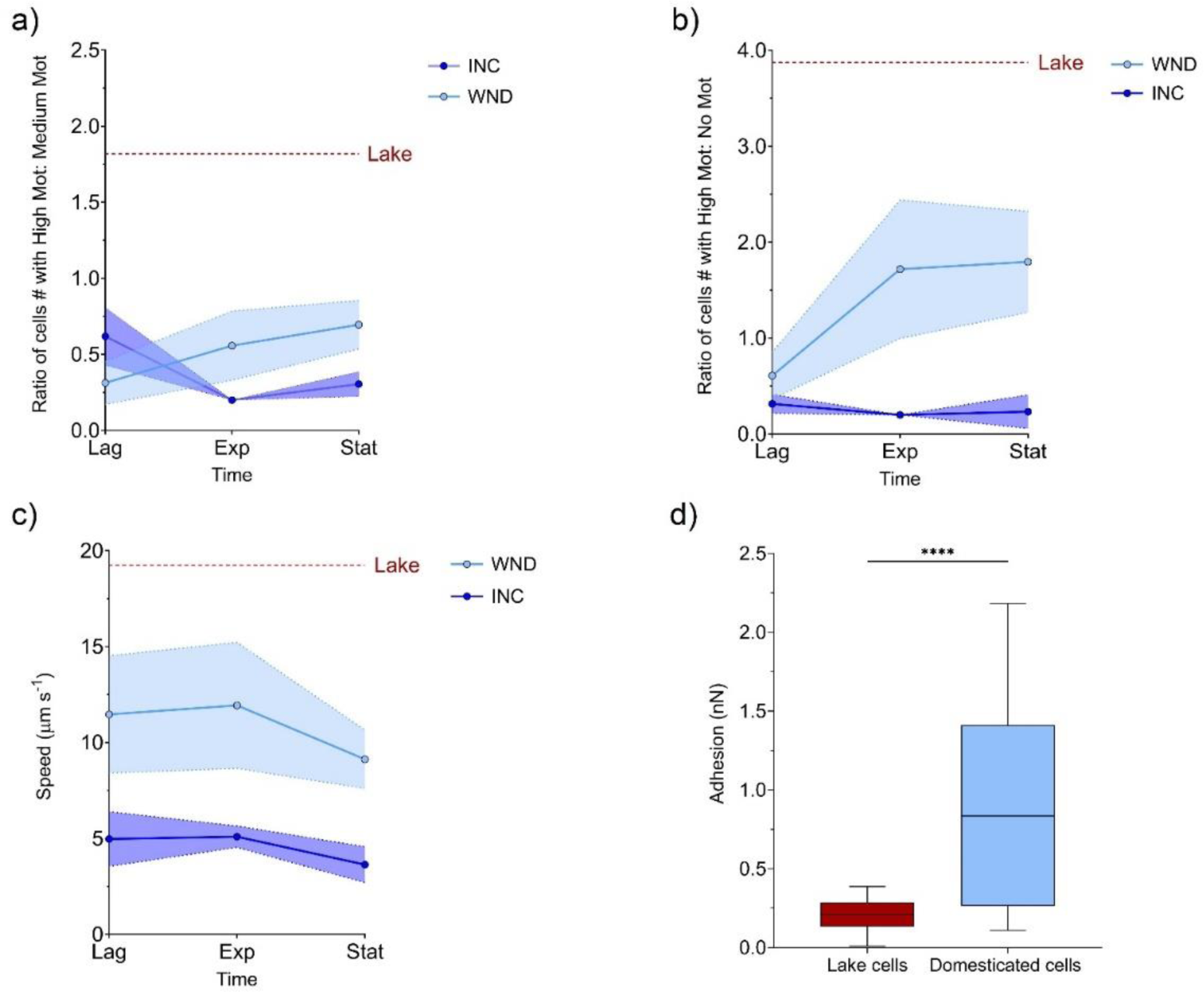
Evolution of cell motility and adhesion over time in laboratory populations. Scatter plots show **a)** the ratio of cells displaying high motility over those with medium motility and **b)** the ratio between cells with high motility over those with no motility along the growth curve. Red dotted lines indicate the same ratios calculated for lake-sampled cells Red dotted line indicates the average speed of lake cells. **c)** Average speed values decrease with time for laboratory cells. Colored regions represent standard deviation (N=3). **d)** *C. okenii* cells show enhanced adhesion after domestication. The boxplots illustrate the adhesion of cells to an agarose surface for lake cells freshly after isolation (maroon-color); and the domesticated cells (light blue, stationary phase). The lab-grown cells show a significantly high adhesion interaction with the surfaces, indicating an increase in biofilm forming ability. Unpaired *t* test, *P* < 0.01; asterisks indicate statistically significant difference.

### Mechanics of cell swimming

In cells sampled from the lake, SGBs tended to accumulate below the cell center of gravity (*C_H_*; Figure 8a and S6). The center of mass of the SGBs, *C_O_*, was located below *C_H_*. Since the position of *C_H_* coincides with the cell’s center of buoyancy, *C_B_*, which overlaps the center of gravity (Figure 8), the accumulation of SGBs in the lower part of the cell made it slightly aft-heavy, the difference between the mass of SGBs in the fore and aft region of the cell being statistically significant (1.46 *vs* 1.26 x 10^-6^ μg, *p* < 0.05). The low value of *L_W_* (distance from the *C_B_*; Table S1) showed that SGBs were mainly scattered near the *C_B_*, resulting in an average *L_W_/a* ratio of 0.039 (± 0.025), which places lake cells close to the boundary of the phase plot where the orientation stability switches (Figure 9a).

**Figure 8.**
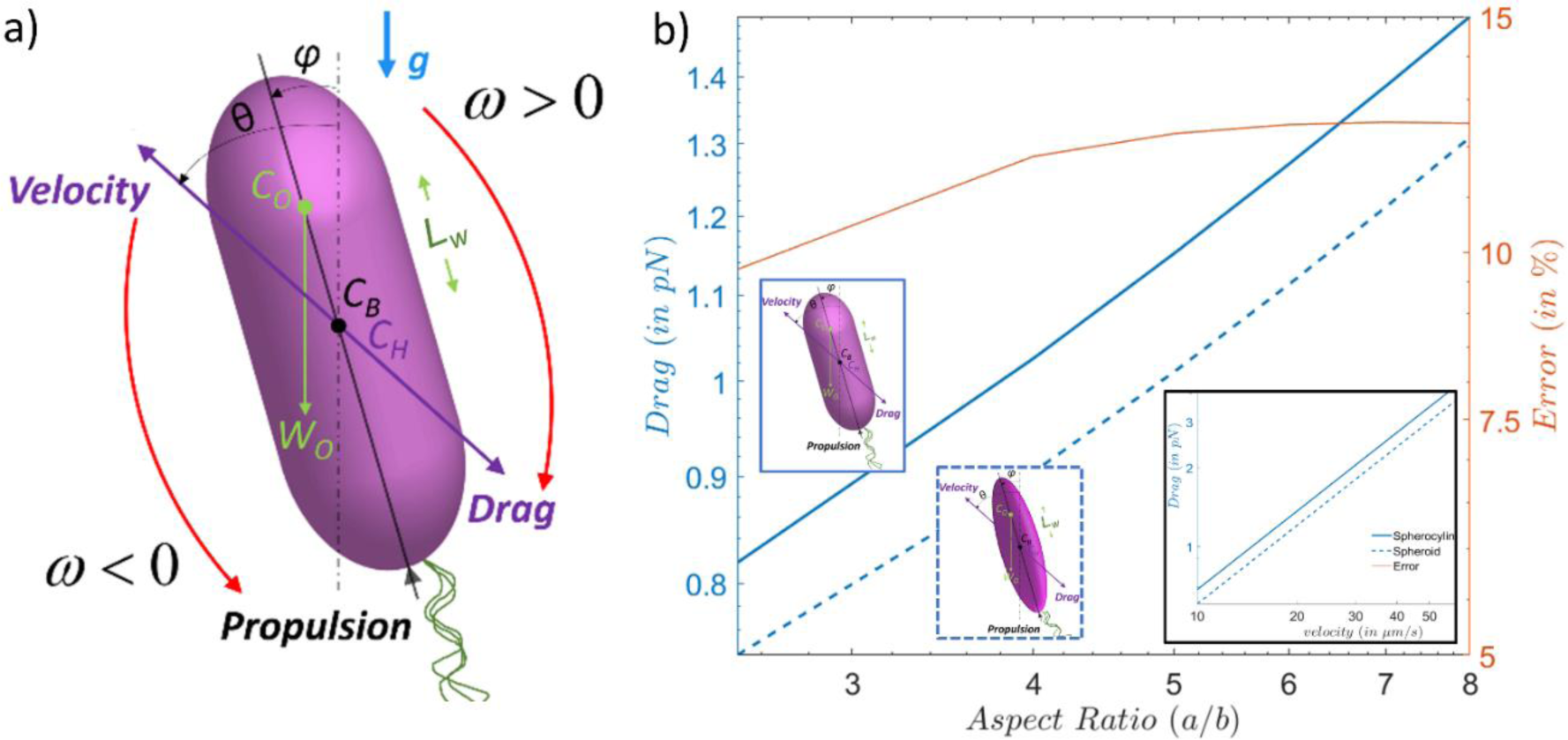
Mechanics of *C. okenii* swimming. **a)** Schematics of the cell-level geometry for the formulation of the reduced-order model. The free-body diagram of all forces and torques (about point *C_B_*) are color marked on the schematics. The swimming of the bacteria cell is considered to be stable when the cell rotates such that its pusher-type propulsion will propel the cell against gravity, *g* (in the above configuration, this is achieved for *ω* > 0). The weight and buoyancy forces act opposite to each other, to give an effective weight, (***ρ***_***cell***_ − ***ρ***_***fluid***_)***Vg***, where *ρ_cell_* and *ρ_fluid_* respectively denote the cell and surrounding fluid densities, *V* is the cell volume, *g* is the acceleration due to gravity (acting downward, in the plane of the figure). **b)** Comparison of drag forces between spherocylinder and spheroid cell geometries for different cell aspect ratios. The y-axis on the right shows the error between the two estimations. For spheroid, the aspect ratio is the ratio between the minor axis and the major axis. For spherocylinders, the aspect ratio is the ratio between the radius of the spherical cap and half of the length of the central cylinder and radius of the spherical caps combined. The maximum error lies below ∼11% for the two values. Alternative calculation of the spherocylinder aspect ratio can yield lesser error values (see Supplementary Text 2 and Figure S7). Inset plot shows the drag force as a function of the swimming velocities.

**Figure 9.**
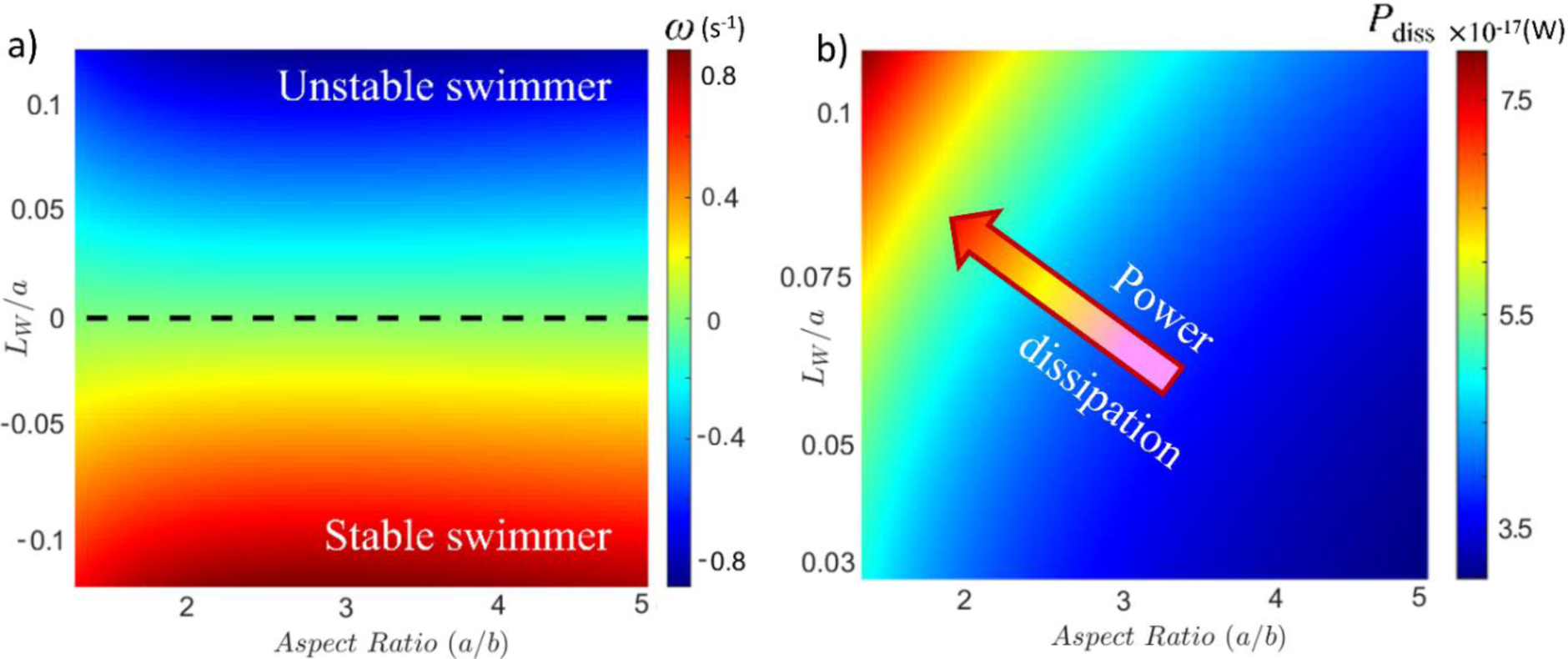
Stability and energetics of *C. okenii* swimming. **a)** Phase plot presents the combined effect of cell aspect ratio (*a*/*b*) and the normalized offset length scale, *L_W_*/*a*, ratio between the position of the cell center of weight (determined by the effective SGBs position) and its major axis. The dashed black line represents the boundary across which the orientation stability switches. Since, only the SGBs position dictates the swimming stability, the aspect ratio does not have any influence on the line of stability. However, the aspect ratio the drag on the cell, thereby determining the rotation rate, i.e., the time taken by the cell to attain equilibrium swimming direction. For a given *L_W_*/*a*, a cell with higher aspect ratio will experience a higher rotation rate, and a higher degree of instability (stability) depending whether it lies above (below) the dashed-black line, respectively. **b)** Power dissipation by swimming *C. okenii* as a function of the aspect ratio and *L_W_/a* values obtained experimentally. For a given aspect ratio, the power dissipated increases with *L_W_/a,* while for a given *L_W_/a*, the dissipated power reduces with aspect ratio.

Conversely, WND and INC cells were characterized by larger offset lengths of 0.91 (± 0.60) and 0.41 (± 0.31) μm (Table S1), respectively, indicating a SGBs distribution shifted to one side of the cell. In fact, although it was not possible to distinguish the fore from the aft section of the cell for WND and INC populations due to the limited motility and the absence of the polar flagellar tuft, we observed marked differences in the globules mass distribution between the two halves of the cell, with 6.0 (± 4.4) and 3.7 (± 2.6) x 10^-6^ μg for WND cells and 2.2 (± 1.6) and 1.2 (± 0.3) x 10^-6^ μg for INC cells. Furthermore, both the increasing size of the SGBs relative to the cell from lag to exponential stage, and the larger offset length *L_W_*, increase the rotational moment that biomechanically influences cell orientation (33).

In addition to the intracellular mass distribution of SGBs, the cell aspect ratio also plays a role in shaping the swimming behavior of *C. okenii*. As shown in Figure 1b, domesticated cells have an overall higher aspect ratio compared to the lake phenotype. This causes WND and INC cells to experience greater drag when moving due to their elongated shape (Figure 9a and S7), making motility even more energetically costly. In contrast, lake cells are characterized by a lower aspect ratio (Figure 1b) and exhibit a spherical geometry, which facilitates swimming as this morphology leads to a reduction in viscous drag (Figures 9a, b and S7).

## Discussion

### Domestication drives changes in phenotypical and intracellular morphological traits

Bacteria are strongly affected by changes in environmental conditions and can modify their morphology in response to environmental cues (26). Several studies have highlighted the synergy between cell shape and motility and this correlation is among the most well-studied morphological relationships (26, 27, 34, 35). In line with these observations, the marked differences we report in cell motility of PSB *C. okenii*, resulting from the adaptation of the cells to the artificial conditions, are accompanied by extensive changes in their morphology. In the two domesticated populations, the difference in volume compared to the lake cells is clear, the latter being significantly larger (Figure 1c).

The reason of such a big difference may lie in the fact that all forms of motility place strong physical and energetic demands on cell shape. For instance, Mitchell (36) calculated that a change in cell diameter of only 0.2 μm can escalate the energy required for chemotaxis by a factor of 10^5^. Cell aspect ratio also seems to play an important role in determining the amount of drag a cell is subject to during motion (37). Our COMSOL numerical simulation is corroborated by the findings presented in other studies (38, 39) where it was demonstrated that drag forces are generally higher for spherocylinders than for spheroid-shaped objects. Overall, modification of cell geometry, together with the distribution of the intracellular SGBs suggests that for the laboratory-grown cultures, motility as a phenotypic trait is energetically expensive, and functionally redundant in the context of the artificial settings. Particularly, the decrease in cell size goes hand in hand with a decrease in the number of intracellular sulfur globules (Figure 1b). The presence of SGBs is known to influence cell volume of PSB (40), as they can comprise up to 34% of their cells dry mass and reach sizes up to 15 μm (41, 42).

Interestingly, the combined increase in SGBs size and aspect ratio observed in the exponential phase of both WND and INC cells suggests that the larger granules size may play a role in the morphological dynamics of cells, also suggested by the higher distance (*L_W_*) of the SGBs center of mass from the cell geometric center. In fact, for elongated cell shapes, the frictional coefficient is higher because the larger surface area increases drag more than the reduction in cross-sectional area reduces it (43). Conversely, the presence of SGBs appears to exert limited influence on motility and cell shape of wild *C. okenii* as, given its much larger average volume, the granules are proportionally smaller compared to domesticated cells (Figure 1f) and located around the cell geometric center (low *L_W_*). The low aspect ratio with respect to WND and INC cells substantiate this hypothesis (Figure 1b). However, despite our observations, a direct relationship between SGBs size and cell shape has not yet been established since, barring a few observations (44, 45), specific cellular localization of SGBs has not been resolved and they appear to be randomly localized in PSB species.

The increase in the aspect ratio of WND and INC cells in the transition from lag to exponential growth phase may also depend on their physiological growth stage (Figure 1b). Commonly, in rod-shaped bacteria, as is *C. okenii*, width remains constant during the cell cycle while length increases exponentially (41). This elongation is needed for the accumulation of FtsZ, a cell division protein that assembles a ring-like structure in the mid-cell region to trigger septation, which is produced at a rate proportional to cell size (42, 43).

Another important morphological difference observed between wild and domesticated cells is the presence of a polar flagellar tuft (Figure 2). *C. okenii* has approximately 40 flagella arranged in a lophotrichous fashion forming a tuft 20 to 30 μm long (21), which is visible as a single filamentous appendage under phase contrast and SEM microscopy (Figure 2a, b left panel). The forward or backward direction of movement is determined by rotating the flagella clockwise or counterclockwise, respectively (21). As flagellar production is an energetically expensive process, motile bacteria growing in culture media may lose motility (9, 25). In fact, rapid growth is prioritized in batch culture, so loss of flagella may be helpful (8, 46). A study of the evolution of the rod-shaped motile bacterium *Myxococcus xanthus* revealed that under low selective pressure, such as in artificial laboratory settings, bacterial motility can deteriorate rapidly (47). In contrast, in a natural environment, the spatially organized habitat and the few resource patches might increase the rate of motility during evolutionary adaptation (48). Using an experimental evolutionary approach, Barreto *et al*. (2) traced the evolutionary trajectory of a naturally occurring isolate of flagellated *Bacillus subtilis* as it adapted to a typical laboratory environment. The authors showed that domestication reduced the swarming motility of *B. subtilis*. In light of our results, we hypothesize that domestication of *C. okenii* to laboratory conditions causes changes in a trait, i.e., the presence of flagella, that instead is important for the fitness of wild populations.

### Specific content of SGBs influences cell buoyant density

Evidence has been provided that the specific amount of various storage compounds accumulated by cells can affect their size (49). Our results show that the variation in the specific content of intracellular sulfur inclusions significantly determine the buoyant density of *C. okenii* cells (Figure 1g). In fact, SGBs accumulation within the cell confinements increases the sedimentation component of swimming, making overcoming gravity more energetically expensive. As the plot in Figure 1g shows, laboratory-grown cells at the exponential phase are heavier than their lake-sampled counterpart when SGBs are present, making swimming even more difficult. That might be the reason behind the loss of motility in laboratory cells, for which active swimming is no longer crucial for survival, as they are provided with abundant light and sulfide.

This hypothesis is further confirmed by the density data of cells without globules (Figure 1g). In the absence of globules, lake cells are heavier (1.98 ± 0.38 x 10^-3^ μg) relative to the laboratory cells (7.34 ± 4.00 x 10^-4^ and 5.49 ± 1.11 x 10^-4^ μg for WND and INC cells, respectively). Such variations in cell buoyancy can have important implications in the natural environment. *C. okenii* frequently forms high cell concentration layers by concerted swimming in the quest for the optimal light and sulfide conditions (50), where it accumulates locally, forming sulfur globules and increasing its mass. As a result, the sedimentation component becomes a very important loss factor in this situation as sinking and the subsequent upward swimming can lead to bioconvection, a phenomenon of macroscopic convective motion of fluids generated by the density gradient (here intended as difference in weight between adjacent layers of water due to the local accumulation of microbial cells) caused by the directional collective swimming of microorganisms (23, 51).

On the contrary, laboratory cells are almost neutrally buoyant, and, even if not actively motile, can float around but as soon as globules are formed, they sediment down. Our specific cell density values fall within the same range as those measured for *Chromatium* spp. by other studies (23, 52, 53). Specific density results from the interplay between multiple cell characteristics, some of which, such as ribosomal material, proteins, and RNA, are specifically tailored to growth rate and are modulated by regulatory processes. However, it has been reported that the influence of growth rate on cell density is relatively modest (increase of about one unit in the second decimal) (54, 55), as volume gains likely offset greater cellular RNA and protein concentrations.

### Adaptation-induced variation of swimming behavior and photosynthetic performance

The loss of motility in laboratory-grown cells is also particularly evident when cultures were exposed to ambient light, in the absence of a localized light source (Figure 4). We compared unoriented, random motility of lake-sampled cells with laboratory cells cultivated in two artificial conditions (window-sill and incubator), each characterized by a different photoperiod and light intensity (Figure 3a). The experiment showed how *C. okenii* swimming activity progressively reduced with the transition from the natural (lake) to the semi-(window-sill) and most artificial (incubator) condition. Concomitantly, we also observed that, in laboratory cells, this decrease in motility (Figure 4c and S4) was accompanied by an increased phototrophic growth (Figure 6) from window-sill to incubator conditions. Thus, cells might reduce motility, an energetically expensive process no longer required when growing in optimal laboratory settings, in favor of photophysiology. This suggests that domestication of the wild phenotype can result in increased fitness in the laboratory at the cost of losing previous traits, such as motility. This process has been observed in a number of well-studied microbial strains, such as *Escherichia coli*, *Bacillus subtilis*, *Caulobacter crescentus*, and *Saccharomyces cerevisiae* (3–6).

In anaerobic phototrophic sulfur bacteria light is the principal factor driving motility and photosynthetic activity (14). Several studies reported that light is a key parameter influencing cell activity in motile microorganisms (51, 56–58). In particular, the length of the photoperiod under which the cells are cultivated plays a major role in shaping growth rate and swimming behavior of phytoplankton and bacteria (57, 59). A recent study investigating the eco-physiological impacts of bioconvection in Lake Cadagno highlighted how the presence and absence of water mixing generated by the swimming activity of *C. okenii* is consequential to the difference in photoperiod length throughout the summer season (Di Nezio *et al*., under review). At the same time, the authors also reported how laboratory cultures of *C. okenii* cells exhibited higher growth rates when cultivated under a 16/8 h than under a 12/12 h photoperiod in a growing chamber (Figure S2). This result agrees with the higher growth rate and swimming speeds we observed in INC compared to WND cells under similar photoperiods (Figure 1a and Figure 7a-c).

In phototrophic sulfur bacteria, however, domestication does not appear to be completely irreversible; a few studies conducted in Lake Cadagno reported that laboratory-grown PSB cells were able to switch back to metabolic rates (CO_2_ fixation and sulfide oxidation) typical of fresh isolates, after an acclimatization period inside dialysis bags in their original environment (16, 17). Overall, our results strongly suggest that growth, and subsequent adaptation, of *C. okenii* to artificial laboratory conditions following propagation from the natural environment results in the modification of important physiological traits, such as cell motility and growth rate.

### Adaptation to artificial settings and variations in phototactic behavior

In the natural environment *C. okenii* exhibits a phototactic behavior (60), as in Lake Cadagno, where its ability to swim upwards towards light (positive phototaxis), combined with negative O_2_ and positive H_2_S chemotaxis, has been linked to the presence of bioconvection. When tested for phototaxis, wild *C. okenii* cells exhibited a considerably higher light-driven motility than the domesticated population (Figure 5). Interestingly, in the wild population, the number of cells showing a swimming speed classified as ‘high’, according to the speed regimes we defined, was significatively larger after 30 min (*t_30_*) of light exposure than after 90 min (*t_90_*) (Figure 5c). Also, the relative number of motile cells was significantly higher at t_30_ (Figure 5d). The reduction in motility observed between *t_30_* and *t_90_* could arise due to the inverse proportion between photopigments content and light intensity in PSB, an acclimatization strategy to protect the cells from photodamage (61, 62). In fact, during the experiment, wild cells were exposed to a light intensity of 14.6 μmol m^-2^ s^-1^ PPFD, nearly five times higher than the light that reaches the chemocline depth in Lake Cadagno (63). Similar behavior and motility reduction was observed when cells were exposed to a point light source of the same wavelength whose intensity increased from 4.4 to 14.6 μmol m^-2^ s^-1^ PPFD (Figure S3 and Supplementary Text 1). A low content of photosynthetic pigments results in a weaker electron flow through the transport chain, which ultimately impacts the cell response to light. In fact, the absence of phototaxis in mutants of *Rhodobacter sphaeroides* and other purple bacteria lacking the photosynthetic reaction center, highlighted the crucial role that photosynthetic electron transfer plays in determining photoresponse (64, 65). In their review on prokaryotic phototaxis, Wilde and Mullineaux also report that, when exposed to different light intensities, PSB cells are able to respond to light only in the range where photosynthesis is not saturated (66).

Conversely, domesticated *C. okenii* underwent a marked reduction in the ability to respond to light cues (Figure 6). This may be due the fact that motility and phototactic sensitivity of PSB show a great degree of variation depending on the light conditions under which the cells are grown and maintained (67). Already in the early 1930s, Schrammek (68) reported that under continuous illumination in a light cabinet, motile PSB cells lost their ability to swim and deposited as a thick red layer inside the culture vial. Years later, Pfennig (60) observed that *Chromatium* spp. cells cultivated inside vials stopped exhibiting their typical phototactic response when exposed to light intensities in the range of 50-100 foot-candles (∼10 - 20 μmol m^-2^ s^-1^ PPFD) for 8 to 10 hours.

Wild *C. okenii* cells exhibit consistency in the main morphological traits across temperature variations (4°C to 20°C) from natural to laboratory environments, as evidenced by aspect ratio and volume measurements (Figure 1h). While temperature, represents a potential limiting factor with influence on all chemical and biochemical activities (69) and phenotypic traits of bacterial cells in general (70), our observations indicate that over the course of their domestication, they successfully maintain their morphology, a key trait that determines mechanics of cell swimming. Taken together, the reduction of motility observed in domesticated *C. okenii* cells may be a consequence of the adaptation to the high light intensity (40 μmol m^-2^ s^-1^ PPFD) in the cultivation chamber. Consequently, the light source used in our experiment (14.6 μmol m^-2^ s^-1^ PPFD) might have been too low to trigger any phototactic movement. Evidence in support of this consideration is provided by a recent study that investigated photosynthetic rates of PSB under different light intensities (17). The authors reported how photosynthetic activity of PSB *C. okenii* and *Thiodictyon syntrophicum*, cultivated under the same laboratory conditions of our experiment, reached its maximum at higher light intensities (> 30 μmol m^-2^ s^-1^ PPFD) than the corresponding wild populations, despite the same nutrient availability regime. Our results clearly show that propagation of *C. okenii* in the laboratory, where the absence of competition and key parameters for growth such as light, nutrients and temperature are stable and not severely limiting, or rapidly fluctuating, as in the natural environment, causes the loss of phototactic behavior, a trait no longer crucial for survival.

### Shifts in motility and enhancement of adhesion promote biofilm lifeform

It is evident from the phase plot (Figure 9a) that a cell will experience the same dead torque rotation for same *L_W_*/a and two different values of aspect ratio (a/b). This is due to the fact that for a given *L_W_*/a as the cellular aspect ratio increases, two competing effects come to play – the high aspect ratio (a) makes the cell hard to achieve a high ω due to viscous resistance, but (b) the cell experiences higher torque. However, for a given aspect ratio, the dead torque rotation ω increases with an increase in the *L_W_*/a ratio. The cost for active rotation of the cell can be denoted by the power dissipation, *P_diss_ = DU+Rωη*. For cells with higher *L_W_*/a ratio or aspect ratio needs higher active torque and hence higher power dissipation to maintain its up-swimming ability. Thus, high *L_W_*/a is associated with lower stability of the cells. Bacterial swimming has been extensively covered using a prolate spheroid structure. Here we analyze the validity of such a consideration using COMSOL simulation with the backdrop that closed analytical Stokes solution for a spherocylinder body is not available in the literature. In Figure 9b, we validate the results comparing the spherocylinder and a spheroid, demonstrating that at low swimming velocities (or low Reynolds numbers, Re), the variations in the drag force (FD) between the two geometries remain very small. The similarity of the solutions can be seen from the right y-plot, that denotes the error between the simulated values. Across the chosen velocity range (with a maximum of 50 μm s^-1^), the error lies below 4%. The error increases with an increase in the swimming velocity. Given that velocity of the present system is up to ∼20 μm s^-1^, the error is contained within values below 1%. The validation of the simulation was done using Stokes flow for a sphere (Figure S8) wherein at low Reynolds numbers, the drag force and Stokes drag coefficient (CD) compare well with the analytical solutions. The simulation for this analysis is accomplished in COMSOL Multiphysics with fine grids near the boundaries and grid independence test performed for the simulation results.

The alteration of motility patterns is accompanied by a significant enhancement of the cell adhesion to surfaces (Figure 7d), thus indicating an overall shift of the population from a planktonic to a sessile lifeform under laboratory conditions. The regulation of adhesive interactions is associated with emergence of biofilm lifeform, and is known to play a central role in transition from planktonic to sessile states in both in both aquatic and terrestrial ecosystems (71). Research on adhesive interaction of *C. okenii* remains largely unexplored, specifically in the context of biofilm formation. Consequently, future studies aimed at understanding quorum sensing – a key mediator of biofilm initiation – could shed light on the molecular facets of motile to sessile transition in *C. okenii*.

## Conclusion

Microorganisms often face significant environmental stress in their natural habitat and must adapt to constantly changing conditions (72, 73). To survive and thrive in such challenging environments, bacteria have evolved a remarkable array of strategies, such as the formation of spores, biofilm production, and the activation of stress response mechanisms under other stressors, including temperature (34), light and nutrient availability (74), and turbulence (33, 75). Anoxygenic phototrophic sulfur bacteria in their natural habitats face various environmental stressors such as oxygen concentration, temperature fluctuations, light intensity changes, nutrient availability shifts. Particularly light and sulfide are the main limiting factors for these bacteria as they are key elements in the anoxygenic photosynthesis. In Lake Cadagno, inhibition of photosynthesis at increased light intensities and light limitation restricts the layer of high photosynthetic activity to a few centimeters around the depth of optimal photosynthesis (74). If the environment lacks sufficient sulfide, these bacteria can experience slower growth rates or may need to switch to alternative sulfur compounds or adapt their metabolic pathways to make the most of the available electron donors, which can be less efficient than using sulfide (11). For motile species, such as *C. okenii*, the ability to move within their environment, allows them to migrate to areas with better light and sulfide conditions. However, motility can also be lost as a response to environmental stressors such as nutrient scarcity (76, 77) and extreme temperatures (78, 79). Bacteria might shed their flagella to conserve energy and resources, prioritizing survival over motility. For instance, Ferreira *et al*. (77) provided evidence that flagellar loss is induced by nutrient depletion, indicating that flagellar shedding is not a stochastic event but rather a purposeful ejection or disassembly mechanism employed to adapt to nutrient limitations. In contrast, under laboratory-based artificial environments, like continuous cultures and bioreactors, conditions of optimal nutrient sources, temperature, and illumination may render certain traits redundant. In this settings, flagellar loss can result from extended cultivation under conditions where motility is unnecessary (2, 80). Sher *et al*. (81) reported that *Campylobacter jejuni*, subjected to successive passages within a nutrient-rich laboratory medium, manifested a progressive loss of flagellar motility. Regardless of the setting, flagellar loss exemplifies the adaptability and selective pressures influencing bacterial behavior and evolution and may precede the transition from a planktonic to a sessile lifestyle in phototrophic bacteria (82). This transition, in combination with enhanced adhesion of cells, signify a pivotal shift in their ecological strategy, involving not only the physical attachment of these microorganisms to surfaces but also profound changes in their metabolic, physiological, and genetic profiles. It typically occurs in response to specific environmental cues and is accompanied by the formation of intricate communities like biofilms (82, 83).

In this paper, we combined microfluidics, microscale imaging and quantitative analysis, to describe the domestication-driven modifications between wild and laboratory-grown cells of PSB *C. okenii*. We first report the key role of the environmental conditions by comparing three different growth settings characterized by an increasing degree of domestication. We observed marked alterations in several phenotypic traits, such as cell shape and volume, growth rate and distribution of SGBs, between lake-sampled and laboratory-grown cells. We uncover synergistic interrelations between the morphological and cellular density changes, which lead to emergence of altered swimming behaviours, i.e., random motility and phototactic response, of *C. okenii,* in terms of both average speed and relative ratio of motile *vs* non-motile cells. Overall, we observed a progressive loss of the ability to swim and respond to external light cues with the increasing degree of domestication. Our results support the generalization that progressive adaptation to a new environment involves changes in phenotypic traits that often reflect in marked differences in metabolic activity between domesticated and wild microbial populations. However, it is often unclear if the changes associated with domestication are synergistic, and the extent to which phenotypic shifts, compared to genetic drifts, are responsible for the adaptive traits. This work highlights that these alterations are synergistic in nature, and may lead to complete shift in lifeform as reported here. Under prolonged phases of domestication, an otherwise motile *C. okenii* species shifts to sessile biofilm state, supported by the loss of flagella and enhancement of surface adhesion. Our results suggests that, at the level of metabolic resource allocation, lab-grown cells may switch off their flagellar building and beating machinery so as to allocate resources for promoting higher adhesion between cells and local surfaces, thereby driving the biofilm lifeform.

## Materials and methods

*Chromatium okenii* strain LaCa were exposed to two distinct growth conditions and compared with *C. okenii* isolated from the bacterial layer of meromictic Lake Cadagno (13 July 2022). We systematically analyzed significant differences in cellular morphology to reveal adaptations to increasingly artificial environments. To gain a thorough understanding of *C. okenii*’s behavioral responses to various environmental conditions, we analyzed the morphology of the cells, quantified the intracellular SGBs, and the determined the positioning of the flagella. This allowed us to evaluate alterations in cell motility and phototactic behaviour.

### *In situ* cell sampling

Sampling season in Lake Cadagno started in June (after ice melt) and ended in October 2022. The *Chromatium okenii* cells used in the present study were collected on 13 July from a platform anchored above the deepest point of the lake (21 m). Water for biological analysis was sampled from the chemocline through a Tygon tube (20 m long, inner diameter 6.5 mm, volume 0.66 L) at a flow rate of 1L min^-1^ using a peristaltic pump (KNF Flodos AG, Sursee, Switzerland). Samples were kept refrigerated as to maintain the temperature at which they were sampled (4°C) and in the dark and analysed for microbiological parameters within 1 hour after sampling.

### Laboratory cell culture

Purple sulfur bacterium *Chromatium okenii* strain LaCa was grown in Pfennig’s medium I (84) prepared in a 2.0 L bottle using a flushing gas composition of 90% N_2_ and 10% CO_2_ according to Widdel and Bak (85) and was reduced by adding a neutralized solution of Na_2_S x 9H_2_O to a concentration of 1.0 mM S^2-^ and then adjusted to a pH of approximately 7.1. Cells were cultured in 100 mL sterile serum bottles. One set of cells was grown by the window-sill at room temperature (∼20°C) and under natural light conditions (November to December 2021, light/dark period of approx. 10/14 h) while a second set of cells was incubated at 20 °C temperature in a diurnal growth chamber (SRI21D-2, Sheldon Manufacturing Inc., Cornelius, OR, USA) under a light/dark photoperiod of 16/8 h and a light intensity of 38.9 μmol m^-2^ s^-1^ PPFD (Photosynthetic Photon Flux Density), within the photosynthetic active radiation range (400 - 700 nm). Cultures used for swimming properties and phenotypic traits quantification experiments were propagated from a 35/40-day old pre-culture (stationary growth stage) to standardize the starting population physiological status. The experiments were carried out within a fixed period of the day (between 08:30 h and 13:00 h) to rule out any potential artefacts due to possible circadian cycles of *C. okenii*. The specific growth rate was calculated as the rate of increase in the cell population per unit of time (hours). To investigate the effect of adaptation to artificial settings, we used *C. okenii* cells sampled from the lake and we compared them under two growth conditions: (i) the artificial condition of the laboratory window-sill under natural light (hereafter WND), and (ii) the artificial setting of the laboratory incubator under artificial light (henceforth INC) (Figure 3a). Figure 3b shows the main cell features used to describe *C. okenii* morphology.

### Flow cytometry

*C. okenii* natural and domesticated cells strain LaCa were monitored by flow cytometry (FCM) measuring chlorophyll-like autofluorescence particle events. Cell counting was performed on a BD Accuri C6 Plus cytometer (Becton Dickinson, San José, CA, USA), as described in Danza (86). PSB *C. okenii* can be distinguished from the other anoxygenic phototrophic sulfur bacteria inhabiting the bacterial layer of Lake Cadagno based on morphological characteristics (86).

### Cell tracking

To quantify *C. okenii* cell motility, movies were recorded at 10 frames per second for 10 s and converted to image sequences. Cell tracking was performed using ImageJ Particle Tracker 2D/3D plug-in. Images were analysed by intensity thresholding to determine cell locations and link their position in subsequent frames, obtaining the coordinates of the cells at each interval. Cell coordinates at each frame where then used to extract single trajectories (Figure 3a) and calculate the swimming speed. Only trajectories lasting longer than

1.5 s were considered for swimming speed analysis. *C. okenii* cell body length was used as a threshold to distinguish motile from non-motile cells. For lake-sampled and laboratory-grown cells body length was set to 10 and 8 mm, respectively. Trajectories with a net displacement between 1 and 12 body lengths (10 - 120 µm) and 0.5 and 4 body lengths (4 - 32 μm) were selected for lake-sampled and laboratory cells, respectively. Cells with lower displacements were considered non-motile. Filtering was performed using custom Python code, written using NumPy library. The final filtered trajectories (*T)* were used to calculate speed at each time interval for each cell and values were averaged to obtain the mean swimming speed. Calculations were performed with custom Python code using NumPy and Pandas library. The swimming speeds (µm s^-1^) of a population were plotted as a distribution using matplotlib module. Cells with speeds less than 1 body length were considered non-motile (*N*). To calculate the ratio of motile to non-motile cells (*R)*, total cell count (*C)* of a population (obtained by counting cells in individual frames and then averaging over all frames) was noted. The number of motile cells (*M*) was then obtained by subtracting *N* from the total number of trajectories (*T*)

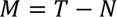

Finally, *R* was calculated as

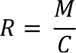

To highlight differences between samples in terms of motility, we arbitrarily defined three different regimes, according to cell swimming speed: no/low motility (< 5 μm s^-1^), medium motility (5 - 20 μm s^-1^), and high motility (> 20 μm s^-1^).

### Volume quantification of intracellular SGBs

To characterize and quantify the biosynthesis and accumulation of sulfur globules (SGBs), cells were sampled from the culture bottles at different time intervals to cover the whole exponential and stationary growth stages. To identify and characterize the accumulation of SGBs in single cells, phase contrast and fluorescence microscopy was carried out, and imaged with high-resolution colour camera. Images were acquired using a Hamamatsu ORCA-Flash camera (1 μm = 10.55 pixels) coupled to an inverted microscope (Olympus CellSense LS-IXplore) with a X100 oil objective. Overall, this gave a resolution of 0.06 μm, allowing us to precisely identify and characterize the SGBs accumulating within single cells. To extract *C. okenii* cell area and SGBs number and dimension (size and volume), pictures and movies of single cells were acquired and analysed as described in Sengupta *et al*. (33, 87).

### Cell morphology and flagellar position

Phase contrast (Zeiss AxioScope A1 epifluorescence microscope) and scanning electron microscopy (Phenom XL G2 Desktop SEM, Thermo Scientific, Waltham, MA, USA) were used to quantify cell morphological characteristics and determine the position of the flagella of *C. okenii*. For SEM imaging, samples were prepared as described in Relucenti *et al*. (88). Cells were sampled from the upper part of the culture vials, to have them as actively motile as possible and exclude nonmotile ones, which sedimented at the bottom. Morphological features, such as aspect ratio and volume, were derived from the contour area extracted by thresholding and ImageJ image analysis. Overall, flagellated cells indicate that they execute pusher type swimming (89).

### Quantification of cellular mass density

To quantify the influence of SGBs on cell density we assumed that the density of structural cell material and the density of the sulfur globules remained constant over the course of the experiment. Other inclusions (i.e., PHB, glycogen) were either undetected or present at a constant quantity and thus considered as components of the cell’s structural material (here cytoplasm). The parameters used in the following calculations are:

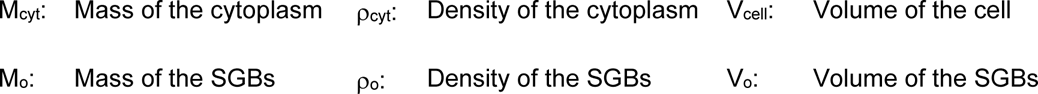

We define total cell mass as:

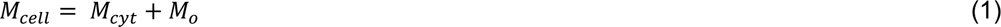

where *M_cyt_* and *M_o_* are the mass of cytoplasm and sulfur globules, respectively. These can be expressed as:

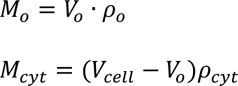

where *V_o_* and *ρ_o_* are the volume and density of SGBs while *ρ_cyt_* is the density of the cytoplasm. Therefore, Equation (1) can be rewritten as:

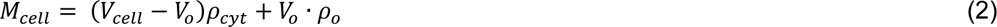

which after simplification, *V_o_* can be converted into:

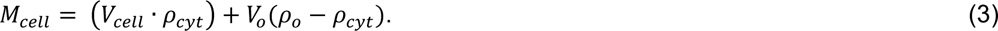

Assuming SGBs to have a spherical shape and cells a spherocylindrical geometry, V_o_ equals 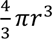 and V_cell_ equals 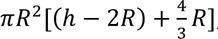, Equation (3) becomes:

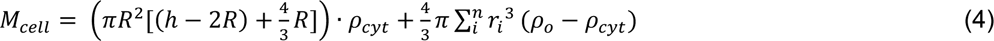

where the summation indicates the sum of the volumes of the *n* SGBs inside a single cell. Dividing Equation (4) by the cell volume, the effective density of the cell can be obtained:

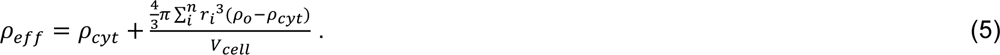

The fraction of the density of the cell accounted for by the SGBs is therefore represented by the term 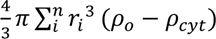.

### Cell phototactic behavior

To investigate the response of *C. okenii* to light, swimming cells sampled from the lake were loaded into rectangular millimetric chambers (microfluidic ChipShop GmBH, Jena, Germany), incubated in the dark for 1 h and then exposed to diffused, low-intensity light from a cold white LED array source (Thorlabs GmBH, Bergkirchen, Germany) placed above the chamber. As a first step, after a 60 min incubation in the dark, one half of the chamber was covered with aluminum foil and the other half was left exposed to light at 14 cm from the LED source, resulting in a light intensity of 14.6 μmol m^-2^ s^-1^ PPFD. Cells were then imaged after 30 (*t_30_*) and 90 (*t_90_*) min for phototactic behavior. Freshly sampled swimming cells kept in the dark were used as a control. Furthermore, the same millimetric chambers were completely covered in aluminum foil, which was then pierced to leave a small circular area exposed to light. The chambers were incubated in the dark for 60 min, and then placed at 14 and 28 cm from the LED source (14.6 and 4.4 μmol m^-2^ s^-1^ PPFD, respectively) and cells imaged after 30 min.

To investigate the potential effects of the domestication process, the same experiment was also performed on laboratory *C. okenii* strain LaCa cells grown in the incubator, the most artificial of the three growing conditions. Cells were grown until their early exponential phase, to have the same physiological growth stage as the wild cells at the time of sampling (90). Laboratory-grown cells in the same growth stage were kept in the dark and used as a control. In both experiments, distribution of cell in the different areas of the millifluidic chamber was determined by ImageJ automatic cell counting on the images obtained at the microscope. Light intensity was measured with a portable LI-180 spectrometer (LI-COR Biosciences, Lincoln, NE).

### Quantification of *C. okenii* adhesion ability

*C. okenii* cells maintained under anaerobic conditions were harvested using a 1 ml syringe equipped with a suitable needle. 0.5 ml of the cell solution was withdrawn from various sections of the cell suspension and subsequently centrifuged at 5000 RPM for 60 seconds. Following centrifugation, 20 µl of the filtrate was carefully transferred onto an agarose gel substrate and allowed to settle onto the substrate for a duration of 10 minutes. Subsequently, a tipless cantilever was calibrated and installed for liquid measurements. For the purpose of force-distance measurements, the cantilever was positioned in an area densely populated with cells. A grid of 20 x 20 measurement points, spaced 1 µm apart, was then recorded (at least 1000 points were measured for each sample) for multiple replicates.

### Modelling mechanics and stability of swimming cells

We developed a cell-level swimming mechanics model to understand the role of the cell morphology and intracellular SGBs, and specifically, delineate the impact of these phenotypic alterations on the orientational stability of the swimming cells (the ability of cells to reorient back to the equilibrium swimming direction after they are perturbed). The model considers different forces and moments acting on a *C. okenii* cell, by virtue of its propulsion, morphology and the SGBs number and intracellular distribution, establishing the factors which determine the cell’s up-swimming stability. A cell generates a pusher-like propulsive force, ***P*** (because of its flagellar dynamics) to maintain its active motion. The weight of the cell (due to combined influence of the SGBs, and the rest of the cell biomass, approximated by cytoplasmic density), and the upthrust on the cell due to the finite cellular volume act in opposite directions. In addition, the cell motion induces a viscous drag (***D***, opposite to the swimming direction) that scales with the cell morphology and swimming speed. Torques on the cell structure are calculated about its centroid (or center of buoyancy, *C_B_*, Figure 8a). The torque contributions on the cell mechanics are the following: effective torque due to the SGBs (when their effective center of mass does not coincide with *C_B_*), the torque originating from the viscous drag (in case of asymmetric cellular geometry), and resistive (viscous) torque due to cell rotation (with rotation speed ω) (33, 87). Based on the physical considerations described in Figure 8a, following equetions emerge from the balance of the forces and the torques:

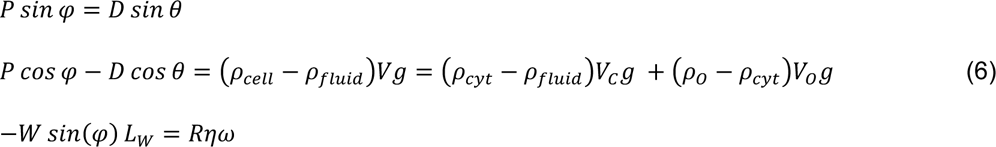

The symbols *ρ, V, W, η* and *L_W_* denotes the density, volume, weight, medium viscosity, and distance from cell centroid respectively. Some of the symbols carry the subscripts *cyt, fluid, C, H, and N* which, respectively, refers to the cytoplasm, surrounding medium (within which the cell swims), the cell, the hydrodynamic center of the cell (which coincides with the cell centroid due to its symmetrical shape), and the SGBs. Density of the cell (ρ_cell_) is given by ρ_fluid_ times sp_cell_, where sp_cell_ is the overall specific gravity of the cell. *φ* is the angle between the line of action of the propulsion force, ***P*** (originating due to the flagellar motion) and the line of action of the gravity vector. Here ω, φ and ***P*** are unknowns, which need to be determined as part of the solution. The motion of the cell does not follow the line of action of ***P***, hence an angular offset θ (an experimentally observable parameter) with the vertical direction is assumed along which the cell moves (Figure 8). φ_N_ is the angle between the direction of the gravity (downward, in the plane of the figure) and the line joining *C*_O_ and *C*_B_ (note φ = φ_N_, since we assume the center of gravity of the organelle to lie on the major axis). *φ*_O_ is the angle between the direction of gravity (vertical line) and the line joining *C*_O_ and *C*_B_ (note *φ* = *φ*_O_, since we assume the center of gravity of the organelle to lie on the major axis). ***D*** denotes the drag force whose knowledge requires the detail of the cellular geometry and its interaction to the surrounding fluid, the details of which are provided below.

Bacteria cells has been traditionally modelled either as a spherocylinder or a spheroid geometry. We have thus simulated the drag for both the configurations and the difference between these values using COMSOL Multiphysics (the validation for the configuration of a sphere is presented in Figure S8). A maximum error of ∼11% is observed between them. Since bacteria are strictly neither spherocylinders nor spheroids, the realistic error should be even less. For the sake of convenient representation without sacrificing the essential physics, we have considered the bacteria as a spheroid shape.

We describe the axisymmetric cell geometry with the generic equation

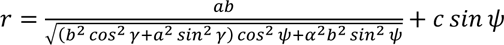

where the symbols 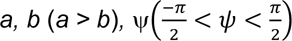, and *γ* (0 < *γ* < 2*π*) represent the major axis length, minor axis length (equal to the semi-major axis length), polar angle, and azimuth angle, respectively. Here *c* implies the deviation from the symmetric shape along the major axis (fore–aft direction) and *r* denotes the position vector of the points on the cell surface (from the origin) as a function of the polar and azimuth angles. With respect to the cell geometry; *a* denotes the full length, and *b* the width of the cell.

The fore-aft asymmetry (value of *c*) is quantified using the phase-contrast microscopy images of the cells whose contours are fitted with Equation (6) and γ = 0, resulting in the form 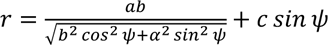. Note that for a symmetric cell geometry (*c* = *0*), the hydrodynamic center (*C*_H_) falls on the cell centroid (*C*_B_), and *L_H_* vanishes. With the consideration that the cell shape may be assumed as a prolate spheroid, the drag of a symmetric prolate ellipsoid is expressed as 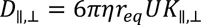 where *U* and *K* are the translational velocity and the shape factor, respectively, while ∥ (⊥) denotes the parallel (perpendicular) direction with respect to the major axis.

The shape factors have the form 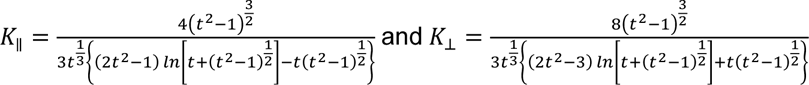 prolate spheroids (91, 92) where *t = a / b*. The net drag on the cell is dictated by its orientation and is given by *D* = *D*_∥_ *cos*(*α*) + *D*_⊥_ *sin*(*α*) (*D*_∥_ and *D*_⊥_ are the drag forces parallel and perpendicular to the major axis of the cell shape, respectively, and α = θ − φ).

*R* represent the coefficient of hydrodynamic rotational resistance and has the form *R* = 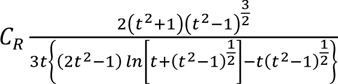 (92) where 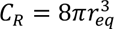. With *R* defined, the viscous torque on a prolate spheroid is estimated using *τ* = *Rƞω* where ω is the angular rotation rate (rad/s). Our aim is to obtain the angular rotation rate **ω** from the above set of three coupled equations (Equation 1). Using the experimentally known values (Table 2), we draw a stability phase-plot (see Figure 9b) that enlists the value of the angular rotation rate as a function of the cell aspect ratio (*a/b*) and the ratio between the position of the cell center of weight (depending on the effective SGBs position) and the length of the long axis (*L_W_/a*). The stability phase plots demarcate the regions of stable up-swimmers from stable down-swimmers, thereby covering a spectrum of swimming stability conditions of *C. okenii* cells representing diverse physiological conditions.

**Table 2.**
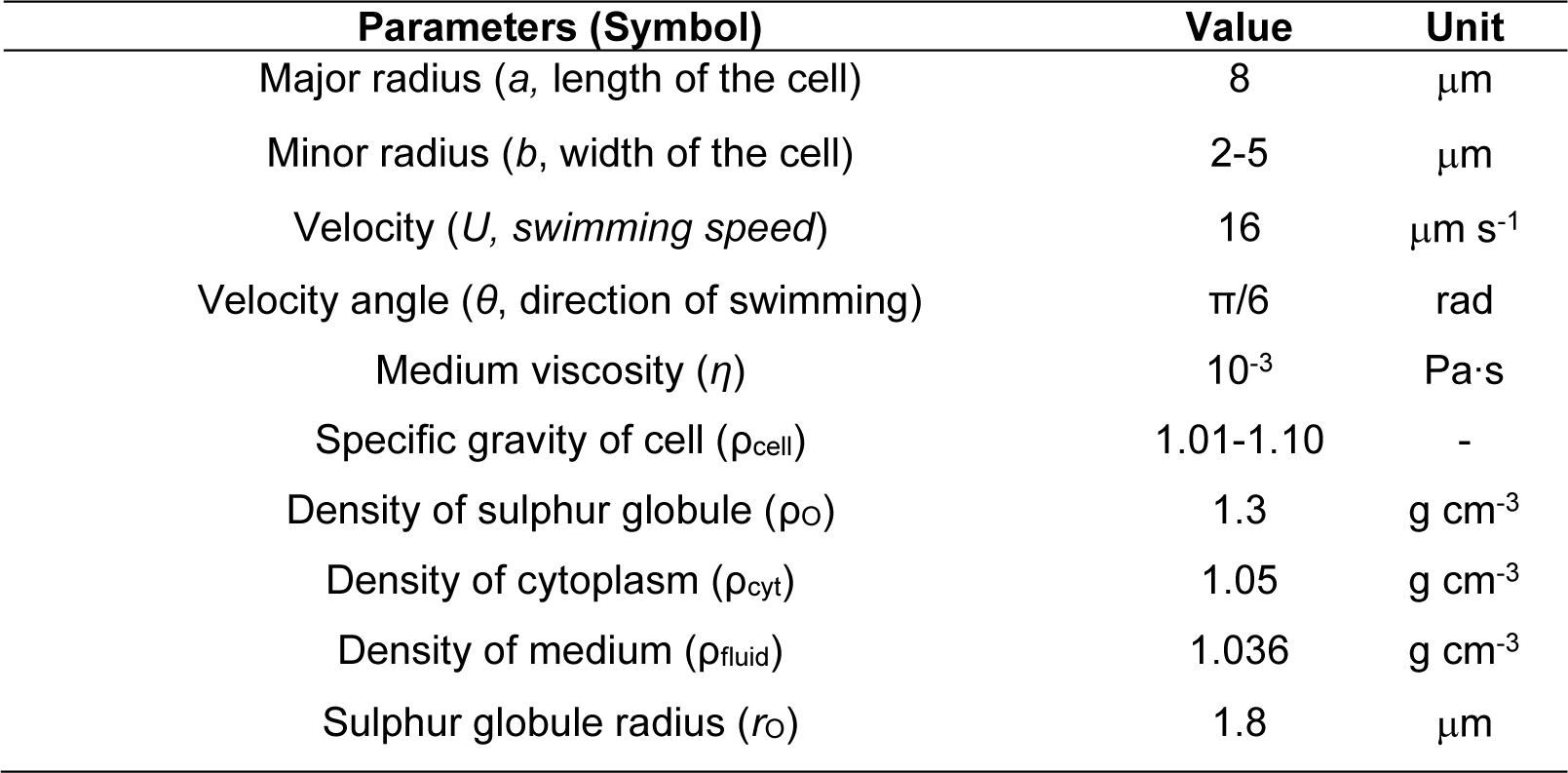
List of parameters used for the computing swimming stability of *C. okenii* cells.

### Statistical analyses

Statistical analyses were performed with GraphPad Prism (version 9 for Windows, GraphPad Software, La Jolla, CA). One-way ANOVA with multiple comparisons using a post-hoc Tukey’s test was performed to compare laboratory-grown *C. okenii*’s cell volume and aspect ratio at different time intervals corresponding to the lag, exponential and stationary growth stages with lake-sampled cells. The same multiple comparison statistical analysis was conducted to compare the number, size, and total volume accumulation of sulfur globules of natural and domesticated cells. Two-way ANOVA with Tukey’s multiple comparisons correction test was used to compare the ratios of motile / nonmotile cells in the phototaxis experiments.

## Author contributions

F.D.N. and A.S. designed research; F.D.N. and S.R. collected samples; F.D.N., I.L.H.O., R.R. and A.S. performed research; F.D.N., R. R., A.G., J.D., and A.S. analyzed data; J.D. and A.S. performed computer simulations. F.D.N. and A.S. wrote the paper with inputs from all authors.

## Data and materials availability

Data and codes are available in the main text, or upon requests to the corresponding author.

### Acknowledgements

We thank Z. Tosheva of the Department of Physics and Materials Science of the University of Luxembourg and P. Principi of the Institute of Microbiology (IM) of the University of Applied Sciences and Arts of Southern Switzerland (SUPSI) for the SEM pictures; S. Forner of the University of Insubria for the experiments using different photoperiods; F. Danza of the Section for Air, Water and Soil Protection (SPAAS) of Canton Ticino for fruitful discussion and constructive comments on the manuscript; the Alpine Biology Centre Foundation (CBA) for laboratory facilities and housing. The authors thank R. Himelrick of University of Luxembourg for helping with the fabrication of experimental chambers and M. Fonseca of SUPSI for supporting with graphics.

