## Supplementary Data for "Synergistic phenotypic shifts during domestication promote plankton-to-biofilm transition in purple sulfur bacterium *Chromatium okenii*"

**This PDF includes:**

Supporting texts 1 & 2

Figures S1 to S8

Table S1

### Supplementary Text 1

Phototactic behaviour was observed in previously dark incubated cells after 30 min of localized LED illumination at two different light intensities (14.6 and 4.4 μmol $m^{-2}$ $s^{-1}$ PPFD; Figure S4). In both light regimes, *C. okenii* showed a larger ratio of highly motile cells in the illuminated sector of the millifluidic chamber, whereas in the shaded one, cells mainly fell into the low- and medium-speed regimes. In particular, the highly motile cells were substantially more when light was lower (4.4 μmol $m^{-2}$ $s^{-1}$ PPFD; Figure S3c), when we also observed a larger fraction of motile *vs* non motile cells (Figure S4c). Analysis of wild cell distribution revealed a significant difference in cell abundance between the illuminated and the shaded region at 4.4 μmol $m^{-2}$ $s^{-1}$ PPFD, while no difference was observed after exposure to higher light intensity (14.6 μmol $m^{-2}$ $s^{-1}$ PPFD; Figure S3b).

### Supplementary Text 2

If we use the spherocylinder aspect ratio as the ratio between the radius of the spherical cap and the half length of the central body cylinder, the error becomes lesser. In this case, however, the total length of the cell for the spheroid and the spherocylinder does not remain the same. For spheroid the cell length is *a* (major axis) while for spherocylinder the cell length is *(a+b)/2*. In that case the error plots are shown in Figure S5.

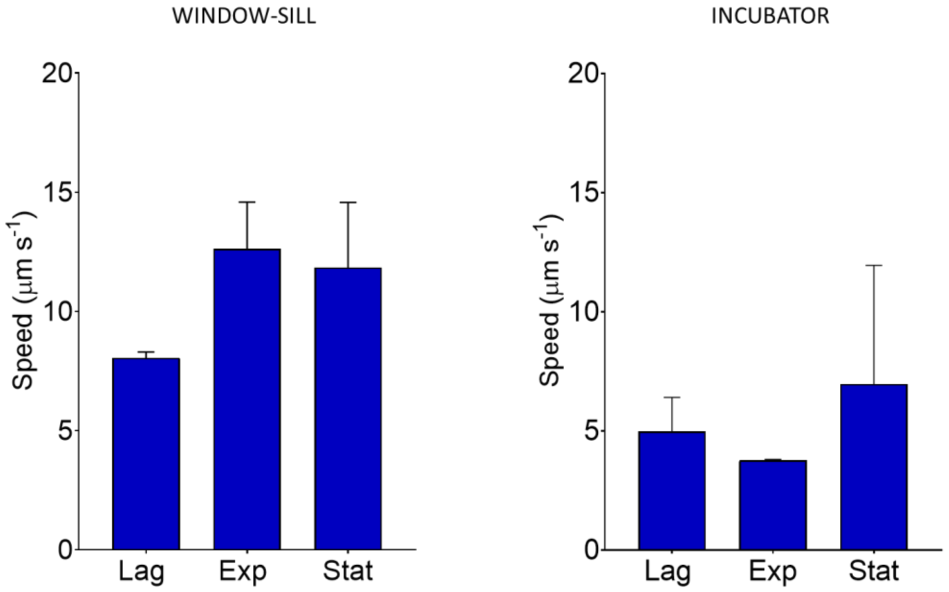


**Figure S1.** Different swimming speeds of laboratory-grown *C. okenii* cells across the main physiological growth stages when cultivated on the window-sill and the incubator. Error bars represent standard deviation (N=3).

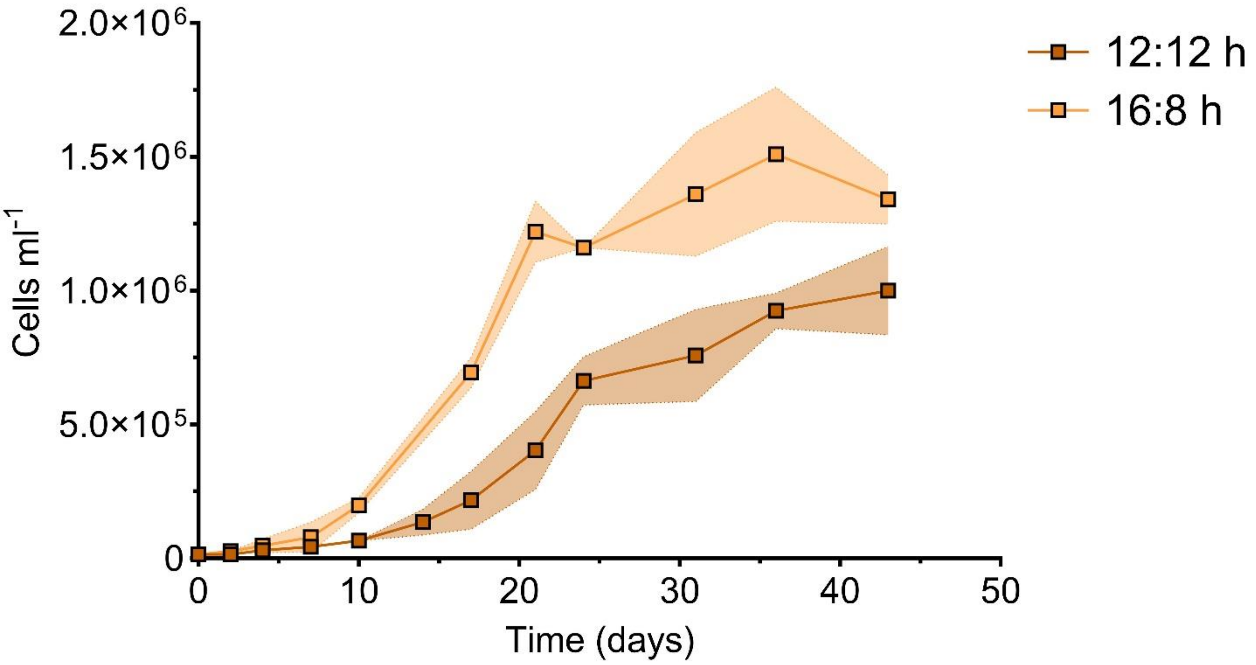


**Figure S2.** Different growth rates observed in WND and INC *C. okenii* cells under light/dark photoperiods of 12/12 h and 16/8 h.

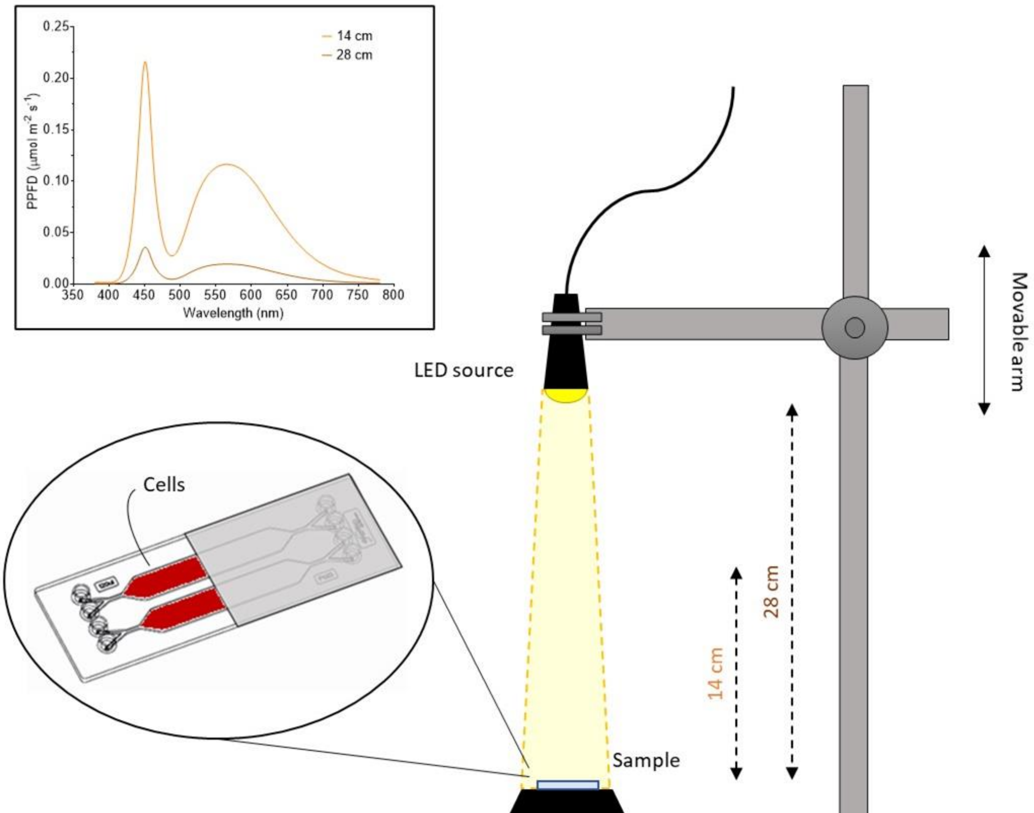


**Figure S3.** Schematic of the microfluidic setup. Inlet shows spectra of the LED light source at 14 and 28 cm distance from the millifluidic chip.

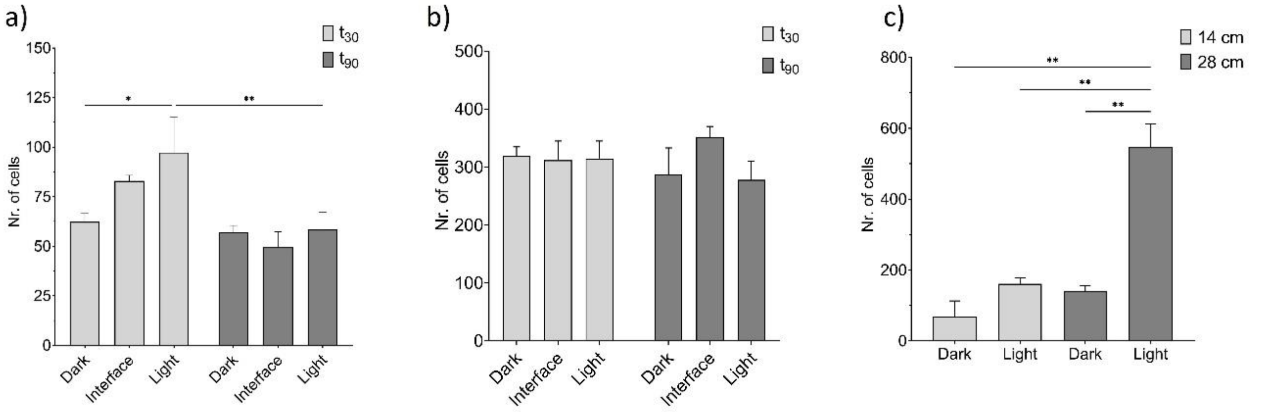


**Figure S4.** Distribution of cells by number in the different sections of the millifluidic device in **a)** lake-sampled and **b)** laboratory-grown cells in the half shaded-half illuminated experiment after 30 and 90 minutes. **c)** Distribution of cells by number in the dark and illuminated areas of the millifluidic device at the two different distances from the point light source. One-way ANOVA, $P < 0.01$; post hoc Dunnet test; asterisks indicate statistically significant difference. Error bars represent standard deviation (N=3).

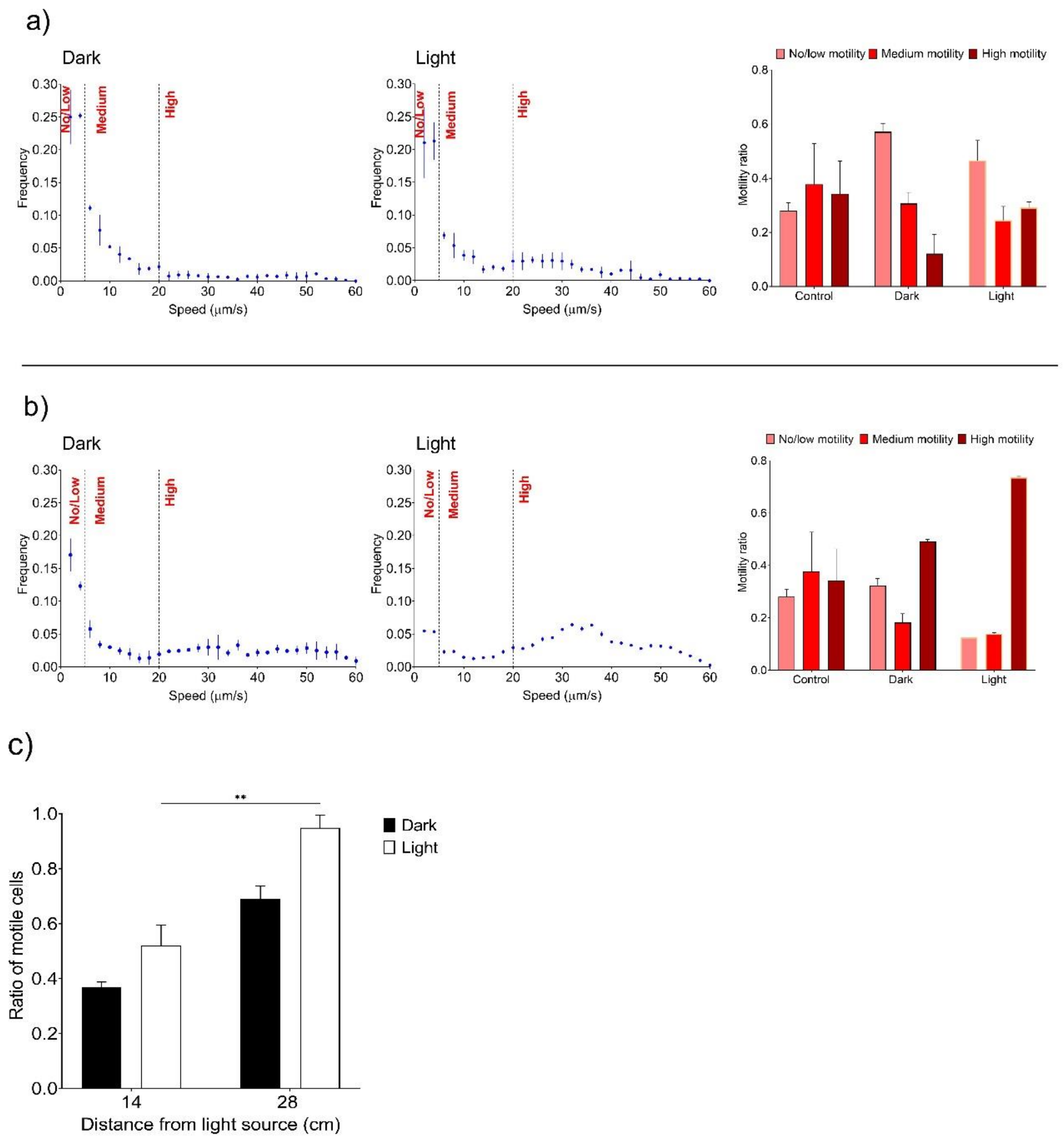


**Figure S5.** Changes in speed distribution of *C. okenii* wild cells between shaded and illuminated sections of a millifluidic device when exposed to **a)** 14.6 and **b)** 4.4 μmol $m^{-2}$ $s^{-1}$ PPFD light intensities. **c)** Different distribution of cells between dark and illuminated regions of the millifluidic device. One-way ANOVA, $P < 0.01$; post hoc Dunnet test; asterisks indicate statistically significant difference. Error bars represent standard deviation (N=3).

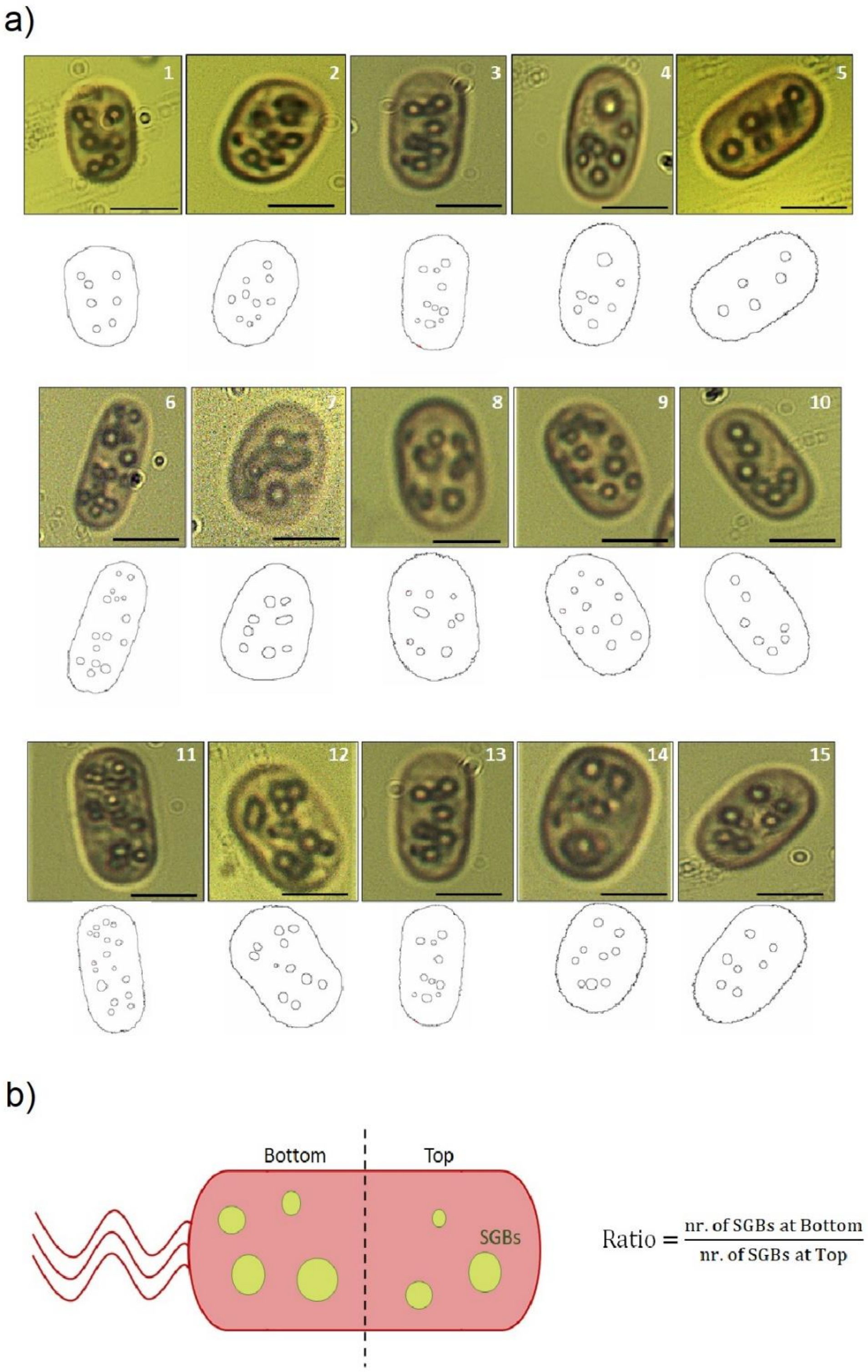


**Figure S6. Determination of sulfur globules position in wild *C. okenii* cells based on single-cell microscopy. a)** Top rows show micrographs obtained by light microscopy with a 100x objective (Methods) of lake-sampled cells. All micrographs are oriented so that the flagella are located in the bottom part of the cell. Image analysis was used to extract the contour of each cell and the position of single sulfur globules in the cell (bottom rows). Scale bar is 5 µm. **b)** Schematic of a flagellated cell showing how the calculations were made.

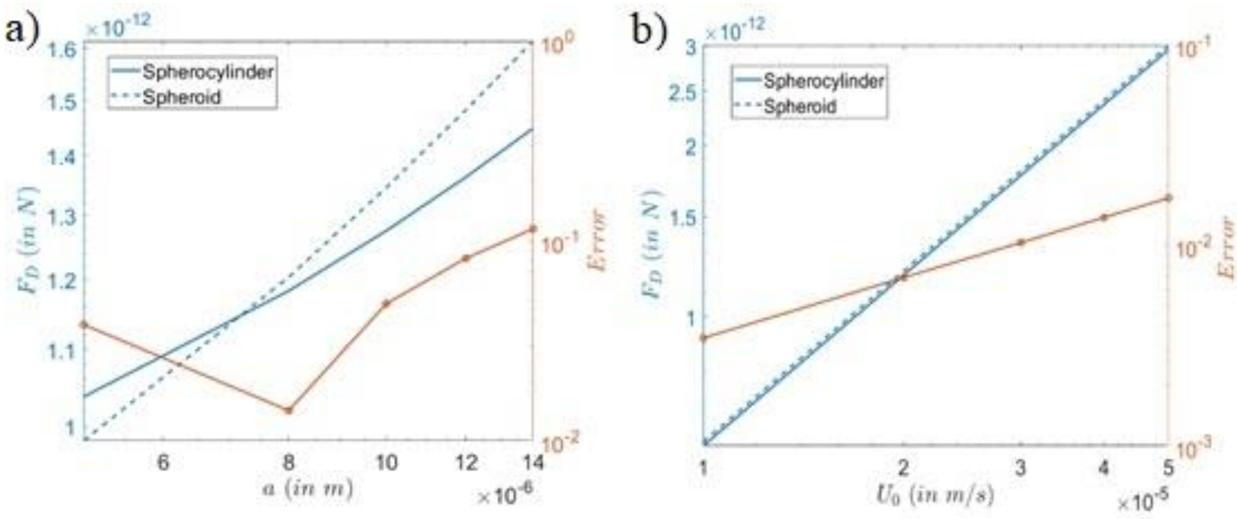


**Figure S7.** Comparison of drag forces between spherocylinder and spheroid cell geometries for different **a)** cell aspect ratios and **b)** swimming velocities. Here the spherocylinder aspect ratio is taken as the ratio between the radius of the spherical cap and the half length of the central body cylinder.

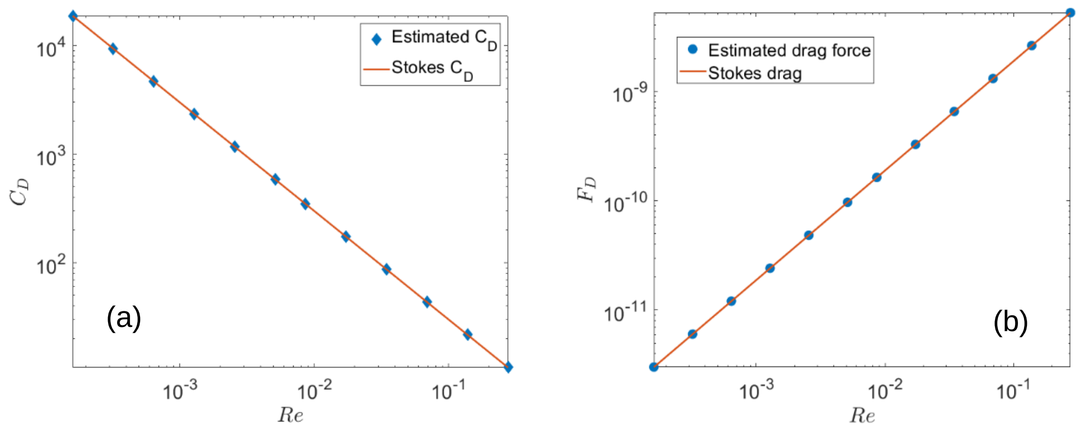


**Figure S8.** Validation of the COMSOL Multiphysics model for estimation of **a)** coefficient of drag and **b)** drag force for spheres.

**Table S1.** Parameters (μm, ± SD) used for modelling mechanics and stability of swimming cells

| | ***a/b*** | $L_w$ | $L_w/a$ |
|---|---|---|---|
| *Lake* | 1.645 (± 0.292) | 0.379 (± 0.270) | 0.039 (± 0.025) |
| *WND* | 2.806 (± 0.866) | 0.912 (± 0.604) | 0.103 (± 0.062) |
| *INC* | 2.231 (± 0.342) | 0.412 (± 0.313) | 0.079 (± 0.063) |
